# Methodologies for Manipulating Cardiomyocyte Physiology: *In Vitro* and *In Vivo* Perspectives

**DOI:** 10.64898/2026.08.11.744256

**Authors:** Run-Zhou Yang, Dan-Hua Liu, Dian-Dian Wang, Sen-Miao Li, Pei-Pei Liu, Shu-Ang Li, Jian-Sheng Kang

**Author notes:** Correspondence should be addressed to Run-Zhou Yang, and Jian-Sheng Kang.

## Abstract

Cardiac tissue is primarily made up of cardiomyocytes, which are regulated by the autonomic nervous system. We have used and developed approaches such as patch clamping and electrical stimulation-combined calcium imaging, computer modeling, optogenetics and chemogenetics combining with video-based Short-Time Fourier transformation (STFT) method to study the physiological activities of cardiomyocytes. The action potential of cardiomyocytes was found to be synchronized with calcium signals, which can be grouped into two categories by STFT. A mathematical model was developed to simulate the changes in electrical activities within cardiomyocytes caused by energy depletion, especially for 2-deoxy-D-glucose (2DG) treatment. Optogenetic and chemogenetics tools, such as ChR2(H134R), OptoXR-β2AR and hM3Dq accelerated beating, while GR, ACR1 and hM4Di inhibited cardiomyocytes’ beating. A video-based STFT method was developed to visualize the beating frequency during these manipulations. An *in vitro* co-culture method was developed to study the relationship between sympathetic neuronal firing and calcium dynamics in cardiomyocytes. *In vivo*, electrocardiograph (ECG) measurements showed that Clozapine N-oxide (CNO) caused heart rates increasement in cTnT-hM3Dq virus injected mouse. However, it had no impact on cTnT-hM4Di virus injected mouse. This study provides comprehensive methodologies for studying cardiomyocyte physiology and manipulating heart rates *in vitro* and *in vivo*.

## 1. Introduction

The heart is among the most metabolically active organs in the body, turning over approximately 6 kg of ATP per day [1]. Cardiac tissue is mainly composed of cardiomyocytes, which can be divided into pacemaker cells and working cells [2]. Pacemaker cells, including the sinus node, atrial internodal tracts, atrioventricular node, bundle of His, and Purkinje fibers, provide spontaneous electrical activity in the heart, while working cells, consisting of atrial and ventricular muscle cells, contract to facilitate the pumping function of the heart [3]. Cardiac action potentials are associated with contraction, a process known as cardiac excitation-contraction coupling [4]. The action potential causes an influx of extracellular calcium, which triggers further calcium release from the sarcoplasmic reticulum, leading to muscle contraction and ATP hydrolysis. Cardiomyocytes have a high demand for ATP due to the repetitive work of the heart muscle. Acute energy deficiency in cardiomyocytes can lead to reduced contractility, arrhythmia and heart attack, while chronic energy deficiency can lead to myocardial remodeling, fibrosis and hypertrophy, which can increase the risk of heart failure [5].

There are several tools and approaches that could regulate cardiomyocyte function, including electrical stimulation, drugs and hormones intervention. Electrical stimulation is a powerful tool in cardiac pacing and defibrillation [6]. Drugs such as β-blockers and calcium channel blockers, are used to regulate heart rate and blood pressure. In addition, the activities of cardiomyocytes can be regulated by the autonomic nervous system, with sympathetic stimulation via hormone such as adrenaline increasing heart rate, myocardial contractility, and conduction velocity, while parasympathetic stimulation reduces myocardial contractility and heart rate [7]. During recent years, optogenetics has emerged as a novel tool for controlling cardiomyocytes. Light-activated chloride pump Natronomonas halorhodopsin (NpHR) and light-sensitive cation channel channelrhodopsin-2 (ChR2) H134R have been expressed in zebrafish and transgenic mice [8] to control heart rate using light and induce disease-like symptoms such as tachycardia, bradycardia, atrioventricular block, and cardiac arrest [9]. Skin-permeable infrared-sensitive rhodopsin ChRmine have been developed for noninvasive optical pacing with a wearable infrared light stimulation vest *in vivo* [10].

Compared to optogenetics, chemogenetics is a genetic engineering technique that enables the modified biomolecules to interact with small molecules, thereby controlling the activity of these biomolecules. Designer receptor exclusively activated by designer drugs (DREADD) technology was developed based on G protein-coupled receptors and modified to be activated by designer drugs such as hM2Di and hM4Di, which are inhibitory receptors, while hM1Dq, hM3Dq, and hM5Dq are excitatory ones [11,12]. Although optogenetics tools have also been applied in manipulating cardiomyocytes, chemogenetics has not been directly employed in controlling cardiomyocyte activities *in vitro* or *in vivo*. Most studies of cardiac chemogenetics involve regulating neurons that associated with cardiovascular system rather than controlling cardiomyocytes themselves. Recently, a synthetic chemogenetic channel PSAM4-5HT3-HC has been developed to modulate the electrophysiology of cardiac tissue [13]. In this paper, we describe the usages of various methods including patch clamping or electrical stimulation-combined calcium imaging, pharmacological treatments and computer modeling, as well as optogenetics and chemogenetics combining with video-based Short-Time Fourier transformation (STFT) method to explore the physiology of cardiomyocytes. These methods constitute a comprehensive toolbox for controlling and regulating the physiological activities of cardiomyocytes *in vitro* and *in vivo*.

## 2. Results

### 2.1 Calcium and voltage dynamics in cardiomyocytes *in vitro*

We utilized patch clamping and simultaneous calcium imaging (Figure 1a-c) to evaluate the traditional cardiac excitation-contraction coupling theory [4]. In cardiomyocytes, intracellular calcium was indicated by Fluo-4 [14], a calcium sensitive dye (Figure 1a, Supplementary movie 1). The action potential and simultaneous calcium signal during continuous beating were demonstrated in Figure 1B. Consistent with the excitation-contraction coupling theory, each action potential of a cardiomyocyte was well synchronized with a calcium spike (Figure 1b), and the peak of the action potential precedes the peak of the calcium signal by approximately 158 ± 40 ms (Figure 1c).

**Figure 1.**
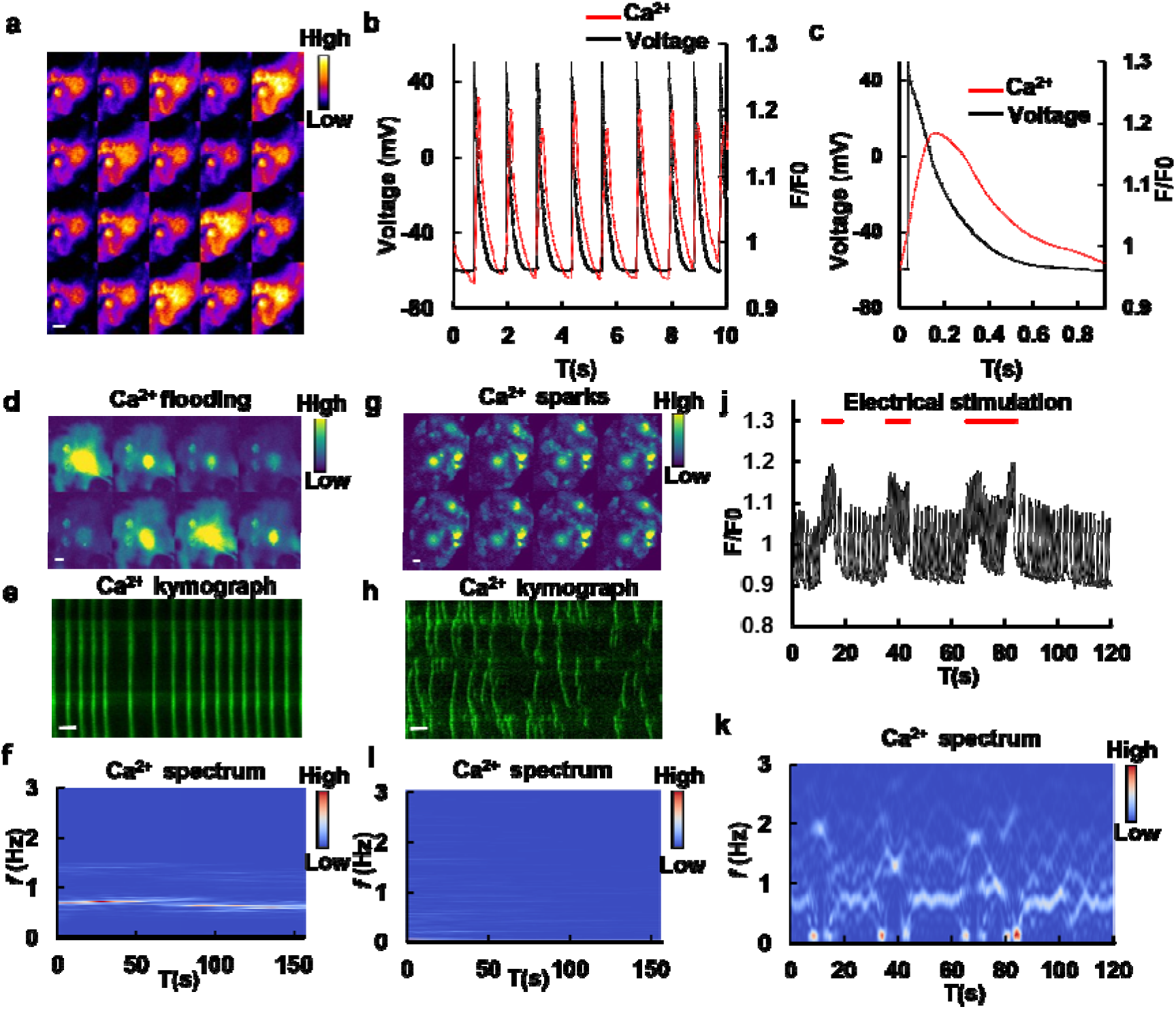
Dynamics of calcium ions in cardiomyocytes. (**a**) Fluorescence images of spontaneous calcium fluctuations in primary cultured cardiomyocytes. Images were acquired at 0.52 s intervals and are arranged from top left to bottom right. Scale bar, 5 μm. (**b**) Simultaneous recordings of action potentials (black) and calcium dynamics (red). (**c**) Representative single action potential curve and simultaneous calcium concentration change. (**d-i**) Patterns of calcium fluctuations in cardiomyocytes could be classified into calcium flooding (**d-f**) and calcium sparks (**g-i**). Representative time-lapse images acquired at 0.13 s intervals, arranged from top left to bottom right. Scale bar, 5 μm. (**e**) Kymograph of Ca^2+^ flooding. Time scale bar, 20 s. (**f**) Short-time Fourier transform (STFT) of signals in Ca^2+^ flooding. Calcium flooding events exhibited a stable frequency. (**h**) Kymograph of Ca^2+^ sparks. Time scale bar, 20s. (**i**) Short-time Fourier transform (STFT) of signals in Ca^2+^ sparks. Calcium sparks were characterized by local and transient fluctuations in calcium concentration without a characteristic frequency. (**j**) Calcium curves of cardiomyocytes in response to external electrical stimulation. The red bars indicate electrical stimulation. (**k**) STFT analysis of curves in (**j**).

We identified two distinct patterns of calcium signals: calcium flooding, characterized by synchronized, cell-wide signals, and calcium sparks, appearing as localized, random events (Figure 1d and 1g, Supplementary movie 2 and 3). Calcium flooding exhibited synchronized signals throughout the cell, while calcium sparks appeared randomly within the cell, as indicated by their kymographs (Figure 1e and 1h). By employing Short-Time Fourier transform (STFT) [15] to calcium signals, these two categories could be distinguished (Figure 1f and 1i). The frequency of signals in calcium flooding remained relatively constant and was equivalent to the beating rate of the cardiomyocytes, whereas there was no consistent frequency for calcium sparks.

To validate the utility of STFT in visualizing beating frequency, we electrically stimulated cardiomyocytes. Each electric stimulation caused additional calcium spikes (Figure 1J, Supplementary movie 4). The spectrogram generated by STFT of calcium signal demonstrated that the frequency of calcium increased upon electric stimulation (Figure 1k). Taken together, the results demonstrated that the action potential and calcium signals were closely related in cardiomyocytes and STFT method could be used to visualize beating frequency.

### 2.2 Calcium and voltage dynamics under energy deprivation

As cardiomyocytes were highly energy consuming during beating, we investigated the effects of energy depletion on calcium and voltage dynamics by using various metabolic inhibitors to target key energy production pathways in primary cultured cardiomyocytes. By visualizing frequency using STFT of calcium signal, inhibiting glycolysis by 2-deoxy-D-glucose (2DG) [16] or oxidative phosphorylation by oligomycin (complex V inhibitor) caused a gradual decrease of the beating frequency in primary cultured cardiomyocytes (first row in Figure 2a and Figure 2b, Supplementary movie 5 and 6). The cardiomyocytes suffered a sudden arrest when applying NaN_3_ (complex III inhibitor) or CCCP (proton uncoupler), characterized by the disappearance of frequency in the spectrogram (second row in Figure 2a and Figure 2b, Supplementary movie 7 and 8).

**Figure 2.**
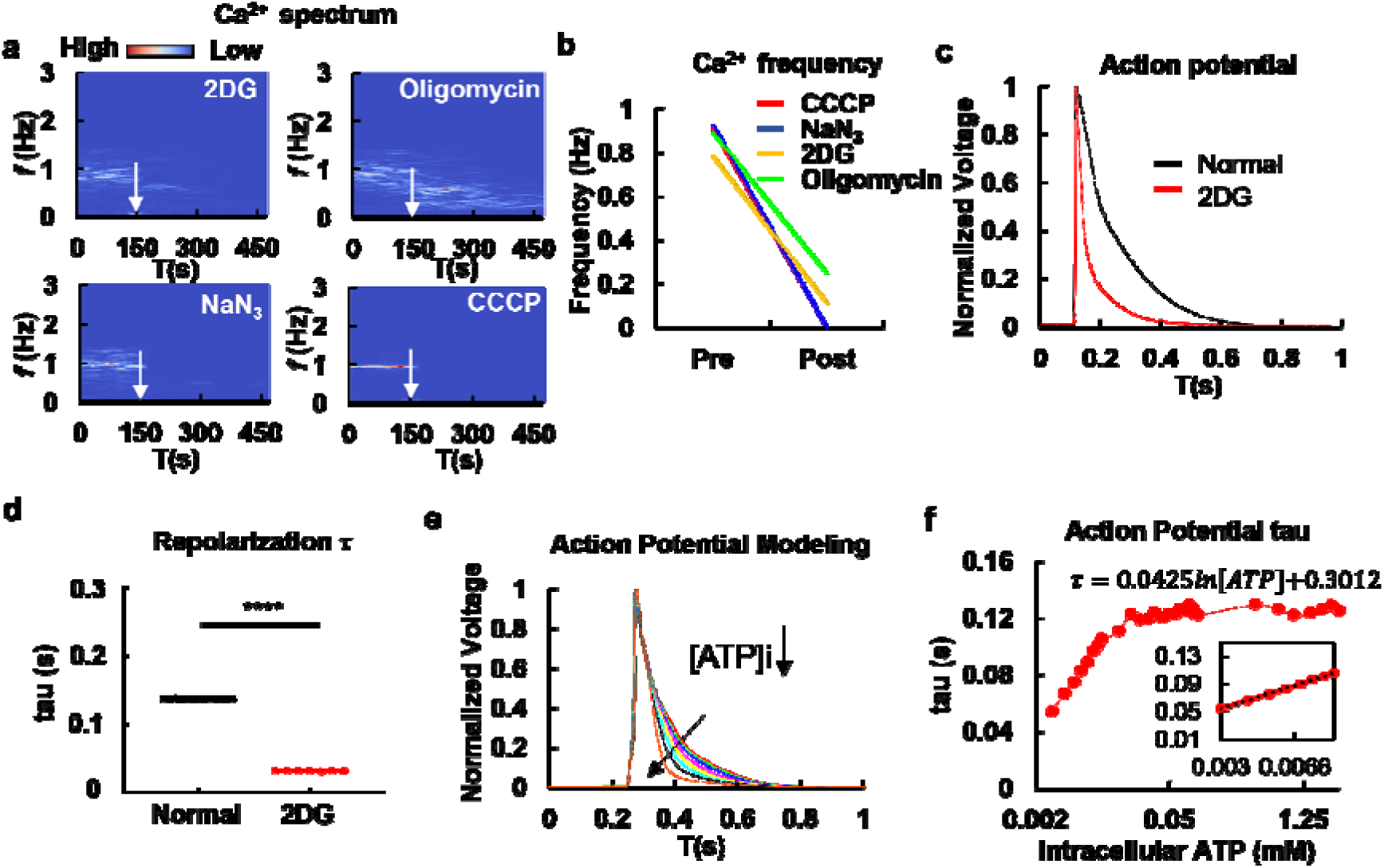
Dynamic changes in calcium and membrane potential in cardiomyocytes during the inhibitions of energy metabolism. (**a**) Short-time Fourier transform (STFT) of calcium signals in cardiomyocytes treated with glycolysis inhibitor (2DG), respiratory chain complex inhibitors (oligomycin, NaN_3_), or an uncoupler (CCCP). The white arrow indicates the time of addition of the inhibitors. (**b**) Frequency quantification of calcium dynamics with drug treatments in (**a**). (**c**) Patch clamp recording of cardiomyocytes membrane potential before (black) and after 2DG treatment (red). (**d**) Comparison of the time constant (τ) in cardiomyocytes before and after 2DG treatment. Single-exponential fitting analysis of the decay phase of action potential reveals a significant difference in the time constant between 2DG-treated cells and control (n = 7 cells, *t-test, p < 0.0001*). (**e**) Computer simulation of action potentials in cardiomyocytes with varying intracellular ATP concentrations ([ATP]_i_). The arrow indicates the direction of decreasing [ATP]_i_ values. (**f**) The relationship between changes in intracellular ATP concentration and decay constants of action potentials. The inset demonstrates that *τ* is linear to the logarithmic curve of ATP concentration within the range of 0.002-0.01 mM. R² = 0.9987.

Interestingly, the treatment with 2DG altered the shape and dynamics of action potentials, resulting in a shorter repolarization period (Figure 2c). Using one-exponential decay fitting, we determined the time constant (τ) of the repolarization phase and found it significantly reduced in 2DG-treated cells (*t-test*, *p* < 0.0001, Figure 2d).

To investigate the underlying mechanisms, we utilized a modified Rasmusson model [17] incorporating the ATP-sensitive potassium channel current (I_KATP_). By decreasing the parameter of intracellular ATP in the model, we found that the repolarization period became shorter, resembling the curve observed in 2DG-treated cells (Figure 2e). We calculated the τ of simulated AP at different [ATP]_i_ (intracellular ATP concentration) levels and plotted the curve of τ versus [ATP]_i_ (Figure 2f). Surprisingly, a linear relationship with a small slope of 0.0425 between τ and logarithmic of [ATP]_i_ was observed and fitted with an R squared value of 0.9998 (Figure 2f). Furthermore, the τ exhibited a stationary value when [ATP]_i_ was more than 1 mM (Figure 2f). Collectively, energy deficiency in cardiomyocytes was detrimental and associated with abnormal electrophysical activities.

### 2.3 Optogenetics in cardiomyocytes

As a methodology of manipulating the physiology of cardiomyocytes, optogenetics tools were applied to primary cultured cardiomyocytes. ChR2(H134R) was expressed in cardiomyocytes and used to stimulate beatings through light stimulation (Figure 3a-d). The continuous beatings of cardiomyocytes were captured by a microscope video camera upon light stimulation (Supplementary movie 9). To quantify beating frequency, we developed a method based on video analysis. Kymographs were generated from bright-field images of beating cardiomyocytes (Figure 3b, top panel), and their intensity was analyzed using Short-Time Fourier Transform (STFT) (Figure 3b, bottom panel). The spectrogram showed that the beating frequency of cardiomyocytes increased dramatically upon blue light stimulation (Figure 3b, bottom panel). Patch-clamp recordings demonstrated that single light pulses elicited single action potentials (Figure 3c). Varying light pulse durations from 20 to 500 ms consistently evoked action potentials (Figure 3d). However, prolonged stimulation with a 500 ms light pulse resulted in a significant refractory period (Figure 3d). This observation likely explains the observed phenomenon of initially regular cardiomyocyte beats transitioning to irregular beating patterns under sustained light stimulation (Figure 3b, bottom panel).

**Figure 3.**
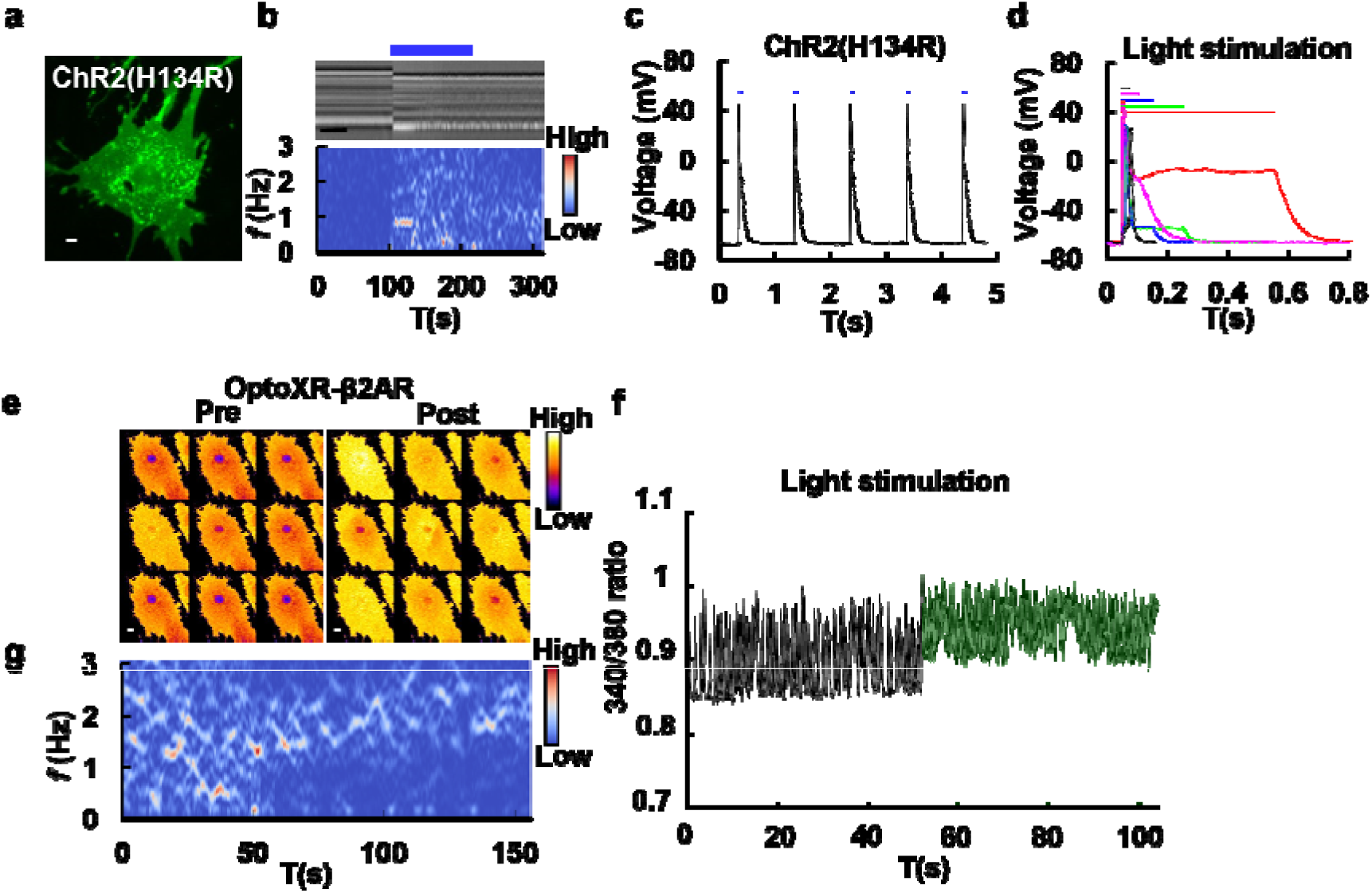
Optogenetic activation of cardiomyocytes. (**a**) Fluorescence image of cardiomyocyte expressing ChR2(H134R)-EGFP, Scale bar, 5 μm. (**b**) Light stimulation induced ChR2(H134R) expressing cardiomyocytes to beat. Kymograph (top) and spectrogram (bottom) based on bright field video demonstrated increased beating frequency of cardiomyocyte after light stimulation. (**c**) Light stimulation could trigger the firing of action potentials in cardiomyocytes expressing ChR2(H134R). The blue bars indicate light stimulation. (**d**) Comparison of action potentials generated at different durations of light stimulation: 20 ms (black), 50 ms (magenta), 100 ms (blue), 200 ms (green), and 500 ms (red) in cardiomyocytes expressing ChR2(H134R). (**e**) Representative calcium images stained by Fura-2 of cardiomyocytes expressing OptoXR-β2AR before (left) and after (right) light stimulation. The ratio between the fluorescence excited at 340 nm and 380 nm was used to represent Ca^2+^ concentration. Images were acquired with a time interval of 0.13 s and represented as pseudo-color. Scale bar, 5 μm. (**f**) Representative Ca^2+^ concentration curve before and after light stimulation in cardiomyocytes with OptoXR-β2AR expression. The green area indicates light stimulation. (**g**) Representative spectrogram of intracellular Ca^2+^ concentration upon light stimulation in cardiomyocyte expressing OptoXR-β2AR. The white arrow indicates the time of light stimulation.

We also explored the use of OptoXR-β2AR, a light-sensitive GPCR that activates the cAMP-PKA pathway[18], to stimulate cardiomyocyte beating. OptoXR-β2AR was delivered to cardiomyocytes by electroporation and green light (532 nm) was used to stimulate the transfected cardiomyocytes. Since OptoXR-β2AR is tagged with EGFP, there is a spectral overlap between the emission of Fluo-4 and EGFP. Therefore, Fura-2, a ratiometric calcium dye with excitation at 340 nm and 380 nm, was used to indicate the calcium dynamics upon light stimulation [14]. The calcium dynamics in cardiomyocytes were present as ratio images of Fura-2 before and after light stimulations in Figure 3e (Supplementary movie 10). The intracellular calcium level was elevated after light stimulation (Figure 3e, 3f). To visualize the frequency change upon light stimulation in OptoXR-β2AR expressed cardiomyocytes, STFT was applied to the Fura-2 ratio. The result indicated that the beating frequency of cardiomyocytes became regular after light stimulation (Figure 3g).

To inhibit the beatings of cardiomyocytes, inhibitory rhodopsins were introduced into cells. Two of the inhibitory rhodopsins, GR [19] and ACR1 [20] were investigated. Cardiomyocytes were transfected with GR or ACR1 by electroporation (Figure 4a and 4d, respectively). The beatings of cardiomyocytes were recorded by a video camera upon light stimulation and STFT analysis. In GR transfected cardiomyocytes, a kymograph was plotted for the bright filed image of beating cardiomyocytes (Figure 4b, top panel and Supplementary movie 11). The spectrogram of cardiomyocytes was generated from STFT of the intensity of the kymograph (Figure 4b, bottom panel). The spectrogram demonstrated that the frequency of cardiomyocytes decreased dramatically upon green light stimulation (Figure 4b, green bar). When the green light was turned off, the cardiomyocytes resumed beating (Figure 4b).

**Figure 4.**
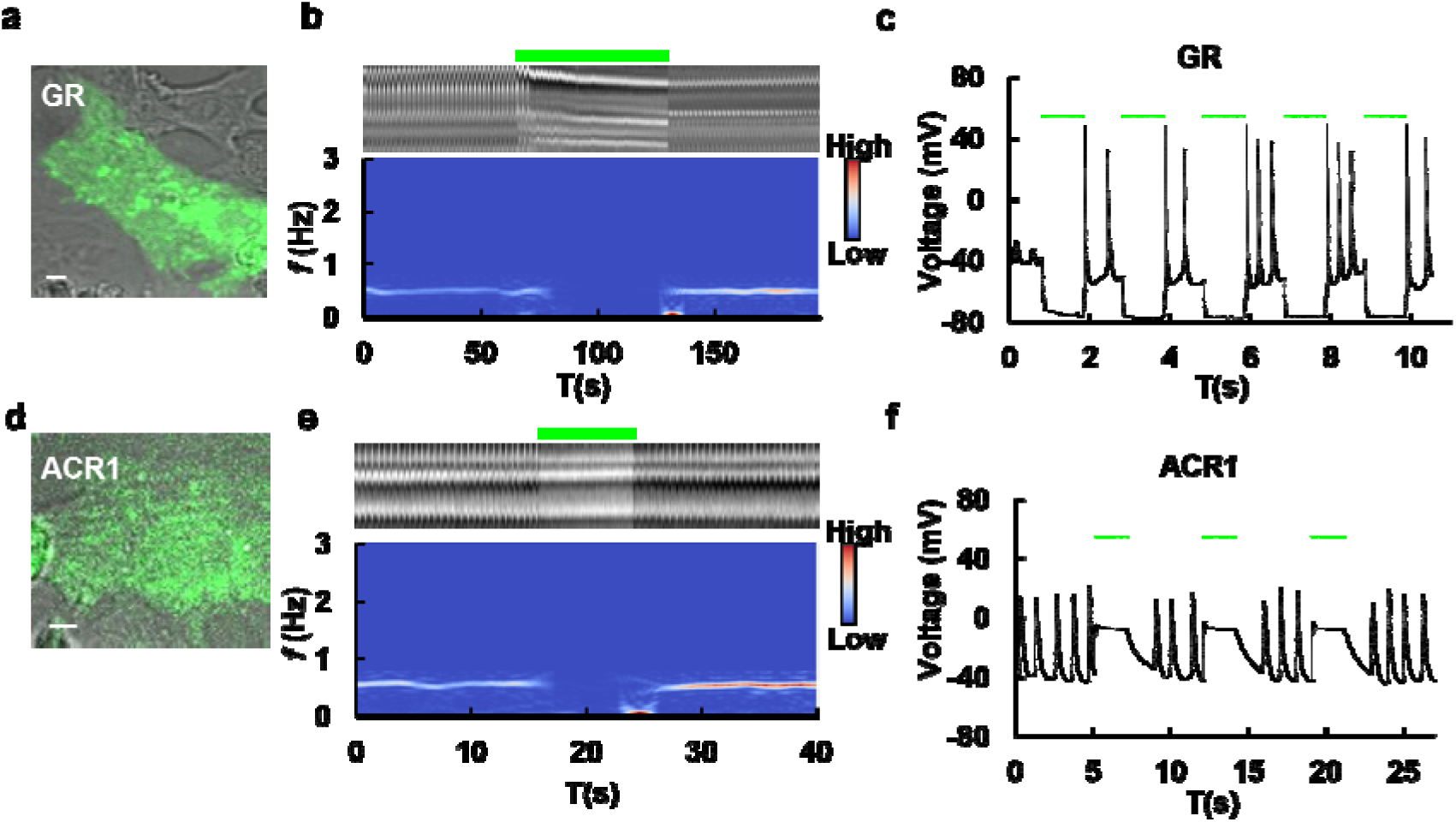
Light inhibitions of cardiomyocytes’ beating. (**a**) Image of cardiomyocytes expressing the proton pumping rhodopsin GR-EGFP. Scale bar, 5 μm. (**b**) Light stimulation could inhibit the beating of cardiomyocytes expressing GR. Kymograph (top) and spectrograms (bottom) based on STFT analysis with bright field video demonstrated decreased beating frequency of cardiomyocytes during light stimulation. The green bar indicates light stimulation. (**c**) Patch clamp recordings of cardiomyocytes expressing GR upon light stimulation. The green bars indicated light stimulation. (**d**) Image of cardiomyocytes expressing the chloride ion channel rhodopsin ACR1-EGFP. Scale bar, 5 μm. (**e**) Light stimulation inhibited the beating of cardiomyocytes expressing ACR1. Kymograph (top) and spectrograms (bottom) based on analysis with bright field video demonstrated decreased beating frequency of cardiomyocytes during light stimulation. (**f**) Membrane potential inhibition in cardiomyocytes expressing ACR1 under light stimulation. The green bars indicated light stimulation. ACR1 activation confined membrane voltage around -20 mV and inhibited action potentials.

Patch clamp recording revealed that the proton pump GR hyperpolarized the membrane potential, leading to inhibited beatings of the cardiomyocytes (Figure 4c). However, prolonged illumination led to sustained contraction and calcium overload (Figure S1). The length of cardiomyocytes transfected with GR was measured before and after light stimulation. The length significantly decreased after light stimulation (*paired t-test*, *p* = 0.006, Figure S1d), suggesting that cardiomyocytes were under sustained contraction. The intracellular calcium indicated by Fura-2 staining demonstrated that a calcium overload happened in GR-transfected cardiomyocytes after light stimulation (Figure S1e-g and Supplementary movie 12).

ACR1, a light gated chloride channel, was also used to inhibit the beatings of cardiomyocytes. In ACR1 transfected cardiomyocytes, a kymograph was plotted for the beating cardiomyocytes in bright filed image (Figure 4e, top panel and Supplementary movie 13). The time-frequency spectrum of cardiomyocytes was generated from STFT of the intensity of the kymograph (Figure 4e, bottom panel). The spectrogram demonstrated the frequency of cardiomyocytes also decreased dramatically upon green light stimulation (Figure 4e, green bar). When the green light was turned off, the cardiomyocytes resumed beating (Figure 4e). Notably, there was a delay between the time switching off the light and the resuming of beatings (Figure 4e). Interestingly, the membrane potential was elevated upon light stimulation (Figure 4f). As a chloride channel, opening the chloride channel brought the membrane voltage closer to the reverse potential of chloride. The delayed recovery from ACR1 inhibition suggested that cardiomyocytes were not efficient to restore intracellular chloride balance when it was disrupted.

As visible light does not penetrate animal skin, light stimulation of heart *in vivo* requires the implantation of LED to the chest, which increases the risk of surgery and infection. Although a wearable vest with infrared light stimulation of ChRmine has been developed to stimulate heart beating [10], there is still a lack of an optogenetic tool that could inhibit the beatings of heart *in vivo*. An auto-illumination system using chemiluminescence was explored here. Gaussia luciferase was fused to the N-terminal of ACR1 and coelenterazine (CTZ) was used to elicit the luminescence (Figure S2a). The luminescence generated by this auto-illumination system was confirmed by a cooled-CCD camera (Figure S2b). As anticipated, adding CTZ inhibited the beating of cardiomyocytes since luminescence activated ACR1(Figure S2c), while the control didn’t show response to CTZ treatment (Figure S2d-e). Collectively, both ChR2(H134R) and OptoXR-β2AR could be used to stimulate cardiomyocyte beating, while GR and ACR1 were used for inhibition. The successful inhibition of cardiomyocyte beating upon CTZ addition in auto-illumination system demonstrated the potential of this system for *in vivo* applications. All the frequency changes can be visualized with STFT of calcium signal or recorded video.

### 2.4 Chemogenetics in cardiomyocytes

Although chemogenetics have been widely applied in neuroscience, seldom has been reported about its application in heart and cardiovascular system. We investigated the application of chemogenetics in cardiomyocytes. The expression of hM3Dq and hM4Di was driven by the cardiac-specific promoter, cTnT, in cardiomyocytes. The cardiomyocytes were transfected by AAV virus expressing cTnT-hM3Dq or cTnT-hM4Di. cTnT-GCAMP7s was co-transfected to monitor the calcium signals in cardiomyocytes. The calcium signals were monitored during Clozapine N-oxide (CNO) application. The addition of CNO revealed that hM3Dq could accelerate cardiomyocyte beating (Figure 5a-c, Supplementary movie 14), while hM4Di inhibited beating (Figure 5d-f, Supplementary movie 15). By applying STFT to the calcium signals of GCAMP7s, the frequency change can be visualized. The results demonstrated that hM3Dq activation caused a gradual increase of beating frequency upon CNO addition. In contrast, hM4Di activation caused rapid decrease of beating frequency especially within 1 min upon CNO application. Interestingly, the cardiomyocytes resumed their beatings 1 min after CNO treatment. To exclude any potential effects of CNO, cardiomyocytes were transfected with cTnT-GCAMP7s and changes in fluorescence were monitored before and after addition of CNO. The results demonstrated that CNO itself had no effects on cardiomyocytes (Figure S3, Supplementary movie 16).

**Figure 5.**
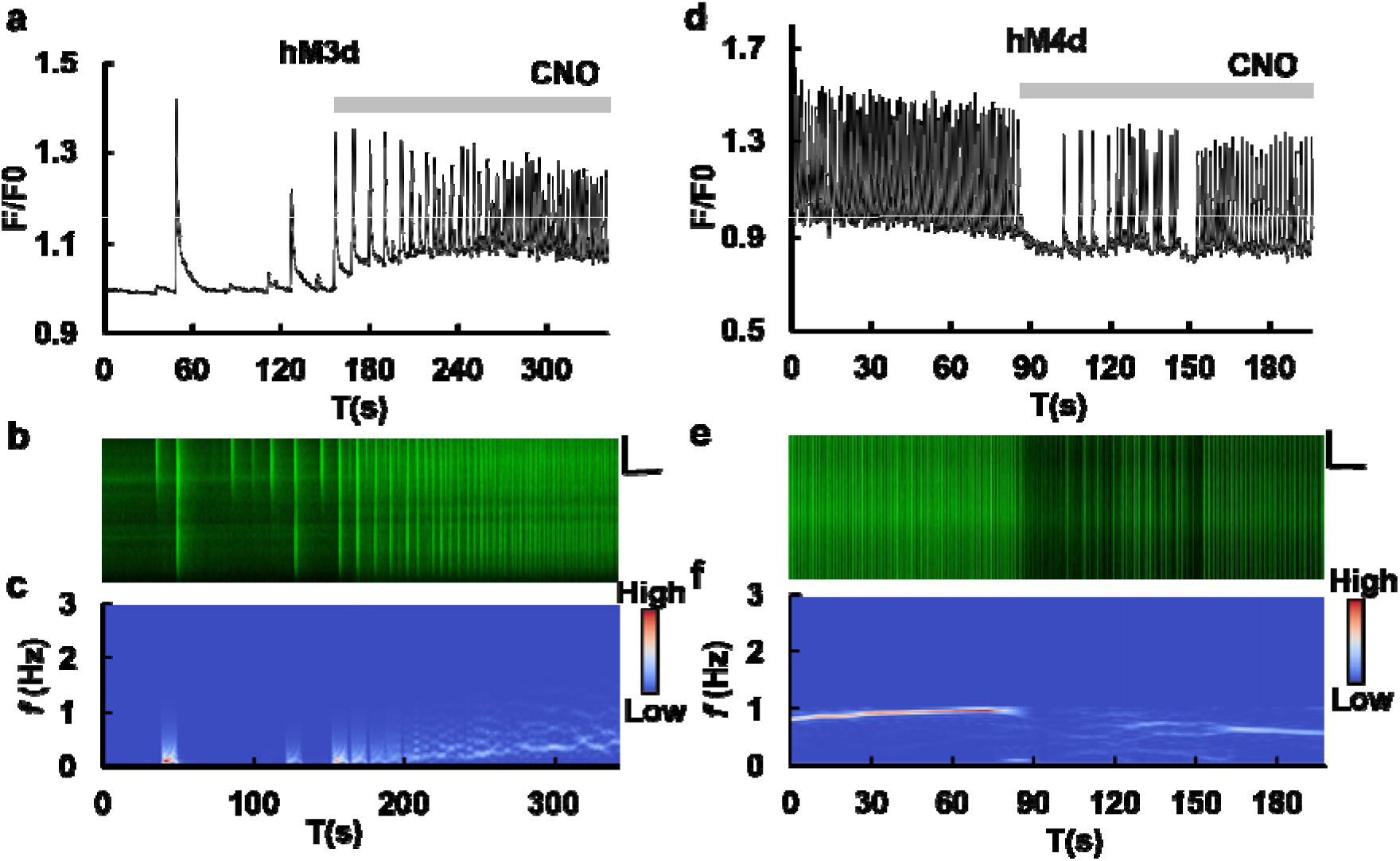
Chemogenetic control of cardiomyocytes’ beating frequency. (**a**) Curve of GCAMP7s fluorescence intensity in cardiomyocytes co-expressing cTNT-hM3d and cTnT-GCAMP7s upon addition of Clozapine-N-Oxide (CNO). The gray bar indicates the CNO application. (**b**) Kymograph of Ca^2+^ in cardiomyocytes co-expressing cTNT-hM3d and cTnT-GCAMP7s upon CNO application. (**c**) Spectrogram of Ca^2+^ in cardiomyocytes co-expressing cTNT-hM3d and cTnT-GCAMP7s upon CNO application. (**d**) Curve of GCAMP7s fluorescence intensity in cardiomyocytes co-expressing cTNT-hM4d and cTnT-GCAMP7s upon addition of CNO. (**e**) Kymograph of Ca^2+^ in cardiomyocytes co-expressing cTNT-hM4d and cTnT-GCAMP7s before and after the addition of CNO. (**f**) Spectrogram of Ca^2+^ in cardiomyocytes co-expressing cTNT-hM4d and cTnT-GCAMP7s upon CNO application.

As the heart is controlled by sympathetic neurons *in vivo*, we further explored the feasibility of chemogenetics in controlling cardiomyocytes by manipulating sympathetic neurons. We established an *in vitro* co-culture system of cardiomyocytes and sympathetic neurons (Figure 6). Briefly, sympathetic neurons were isolated from the spinal cord and co-cultured with isolated primary cardiomyocytes. The human synapsin (hSyn) promoter driven hM3d-mCherry was specifically expressed in neurons, while cTNT promoter driven GCaMP7s was specifically expressed in cardiomyocytes using AAV viruses (Figure 6b). Since the hSyn promoter allows exclusive expression in neurons, calcium signal changes in cardiomyocytes upon CNO addition were attributed to interactions between neurons and cardiomyocytes. In co-cultures with sympathetic neurons, calcium levels of cardiomyocytes dramatically increased upon CNO application (Figure 6c, 6d and Supplementary movie 17). In contrast, calcium levels remained unchanged in cardiomyocytes without sympathetic neurons (Figure 6e-g, and Supplementary movie 18). This method allows us to investigate the interactions between neuron and cardiomyocytes *in vitro*.

**Figure 6.**
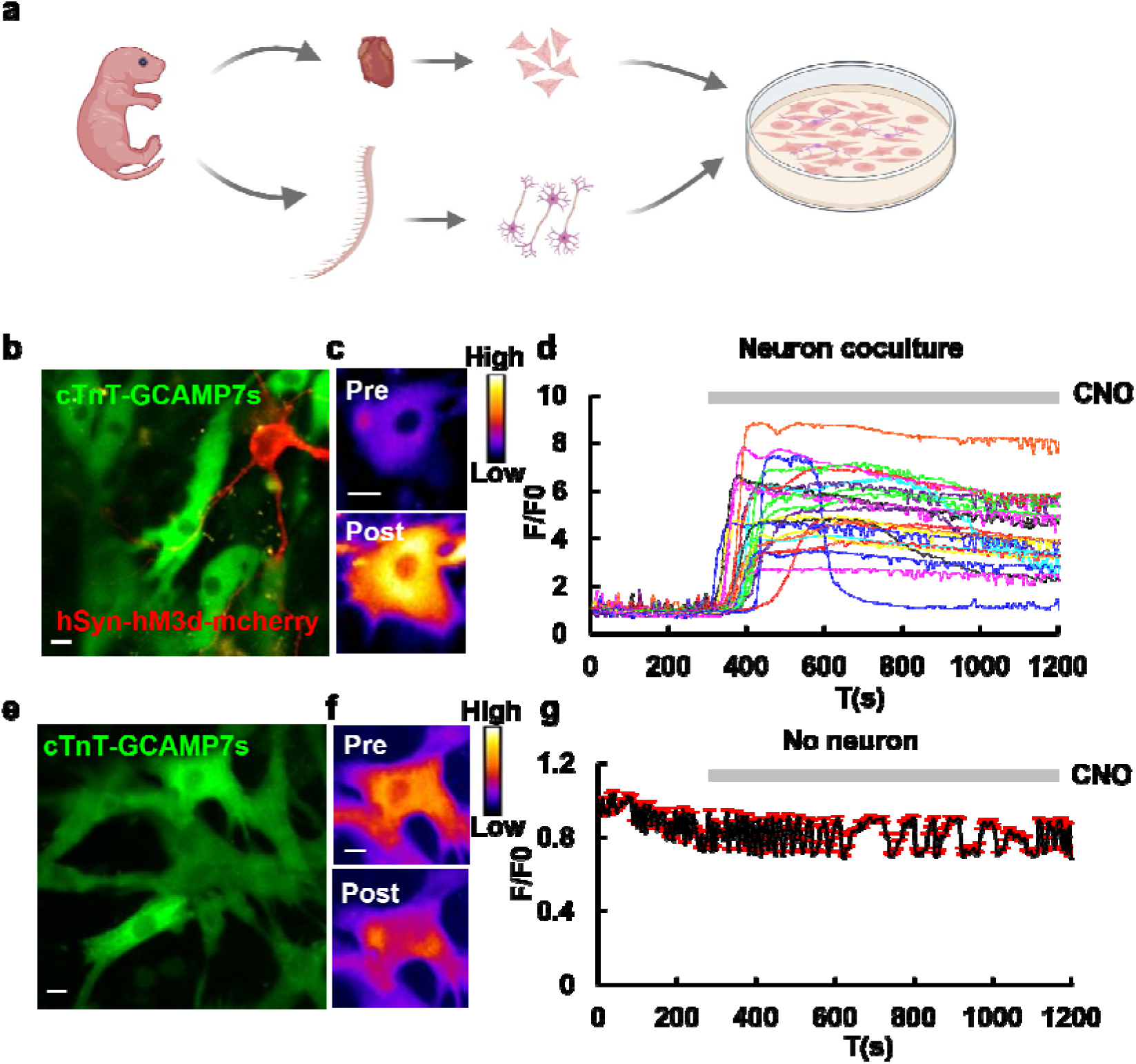
Neuronal control of cardiomyocytes’ activities in co-culture. (**a**) A diagram illustrating the coculture protocol for cardiomyocytes and neurons. (**b**) Fluorescence image of co-culture of isolated cardiomyocytes and neurons, with hSyn-hM3d-mCherry expressed in the neurons (red) and cTnT-GCAMP7s in the cardiomyocytes (green). Scale bar, 5 μm. (**c**) Fluorescence images of cardiomyocytes expressing GCAMP7s before (top) and after (bottom) the addition of CNO in cardiomyocytes and neuron co-culture. Images were represented as pseudo-color. Scale bar, 5 μm. (**d**) Calcium curve of cardiomyocytes in co-culture upon CNO application. Activation of hM3d positive sympathetic neurons with CNO led to an increase of intracellular Ca^2+^ in the cardiomyocytes. The gray bar indicates the application of CNO. **(e)** Fluorescence image of isolated cardiomyocytes expressing cTnT-GCAMP7s without co-culturing with neurons. Scale bar, 5 μm. (**f**) Fluorescence images of cardiomyocytes expressing GCAMP7s before (top) and after (bottom) the addition of CNO. Images were represented as pseudo-color. Scale bar, 5 μm. (**g**) Averaged calcium curve of cardiomyocytes cultured without neurons upon the treatment of CNO (n = 13 cells). The gray bar indicates the application of CNO.

### 2.5 Chemogenetics in heart beating regulation

For an *in vivo* application, we injected cTnT-hM3Dq and cTnT-hM4Di virus into the angular vein of mouse (Figure 7a). Following four weeks after viral infection to allow a sufficient expression level of hM3Dq or hM4Di, the animals were anesthetized, and ECG measurements were performed during the injection of CNO (Figure 7). The ECG signals were subjected to STFT to visualize the change of frequency change. The ECG curves revealed that heart beating rates in mice injected with hM3Dq increased after CNO application (Figure 7b). Following by injection with CNO, there was a gradual increase in ECG frequency (Figure 7c). In contrast, in hM4Di virus injected mouse, the heart beating rates remained unaltered after CNO injection (Figure 7d, 7e).

**Figure 7.**
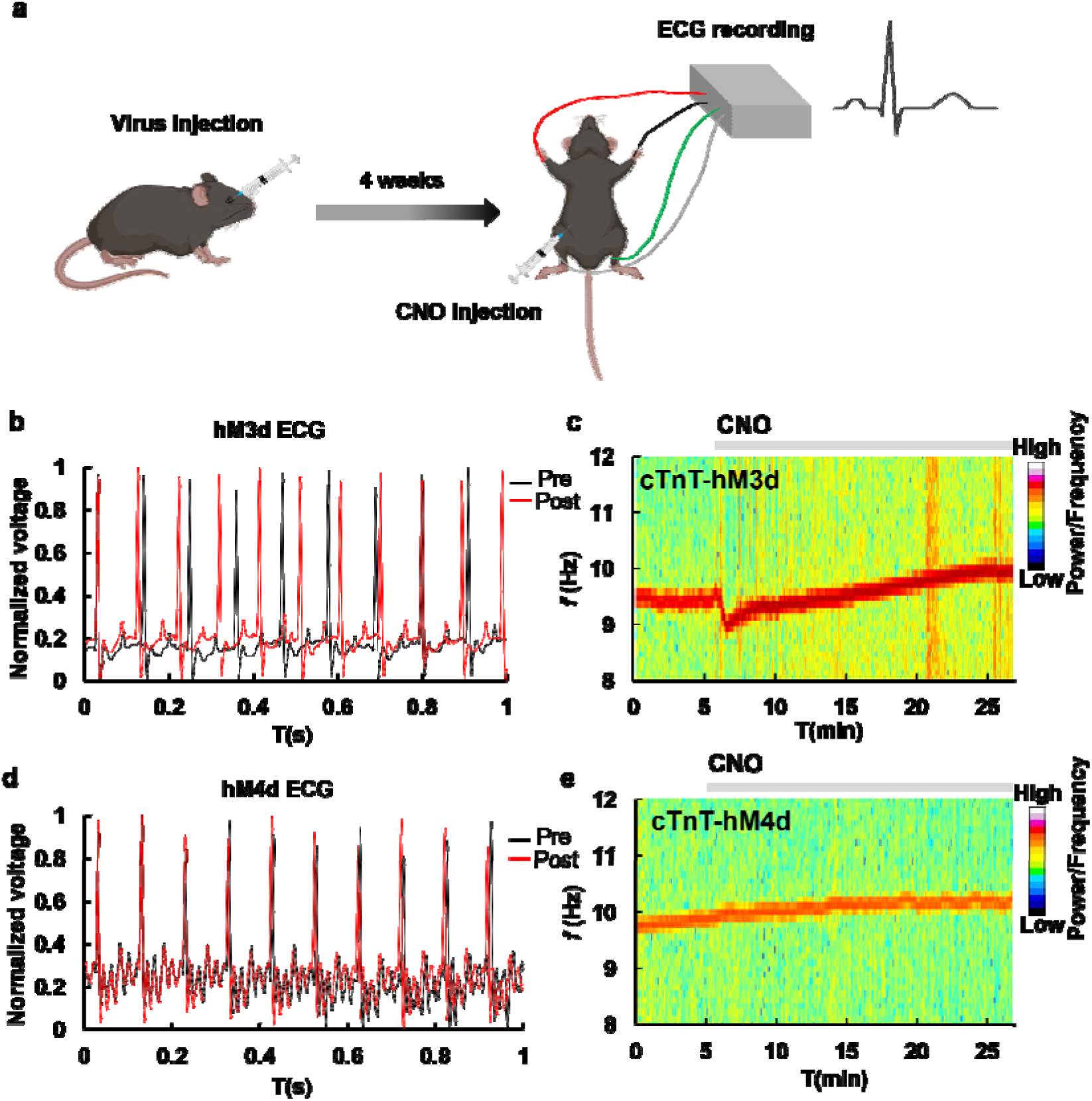
Chemogenetic control of heart’s beating frequency *in vivo*. (**a**) A diagram demonstrating AAV virus injection and ECG measurements. ECG measurements were obtained from mice 4 weeks after angular vein injection with AAV virus expressing cTnT-hM3d. Changes in ECG were monitored during CNO injection. (**b**) ECG recordings in mouse expressing cTnT-hM3d. Representative ECG curves before (black) and after (red) CNO injection were demonstrated. (**c**) Spectrogram of ECG in cTnT-hM3d expressing mouse upon CNO application. The gray bar indicates the application of CNO. (**d**) ECG recordings in mouse expressing cTnT-hM4d. Representative ECG curves before (black) and after (red) CNO injection were demonstrated. (**e**) Spectrogram of ECG in mice expressing cTnT-hM4d upon CNO application. The gray bar indicates the application of CNO.

## 3. Discussion

Cardiomyocytes, the specialized cells comprising the heart’s muscular tissue, exhibit unique structural and functional characteristics. Understanding and regulating the physiology of cardiomyocytes are essential for the development of effective treatments for various cardiovascular diseases. Here we presented applications of various methods for controlling the physiological activities of cardiomyocytes *in vitro* and *in vivo*.

Short-Time Fourier Transform (STFT) is a time-frequency analysis method that is particularly suited for non-stationary signals. It has been widely used in signal processing, audio analysis and image processing. However, it is not suitable for signals with rapid transient changes, varying change rates or low signal-to-noise (SNR) ratios. The continuous beating of cardiomyocytes, exhibiting non-stationary and high SNR ratio characteristics, is well-suited for STFT analysis to extract the frequency. The STFT can be applied to both the calcium signals and bright field videos of cardiomyocytes. A spectrogram exhibiting a single dominant frequency component suggests regular beating of the cardiomyocytes. Conversely, the presence of multiple frequency components within the spectrogram indicates irregular beating. This is beneficial in detection of arrhythmias and monitoring the effectiveness of treatments for arrhythmias.

The effects of energy depletion on calcium and voltage dynamics revealed a relationship between the intracellular ATP and repolarization time constant. Previous studies showed than an outward current increases when cardiac cells subjected hypoxia, while decreased upon intracellular injection of ATP [21]. The ATP sensitive channel is K^+^ specific which were depressed by intracellular ATP at levels greater than 1 mM [21]. 2DG, which leads to ATP decreasing, caused a more rapid repolarization as observed in our study. This indicated that 2DG treatment could lead to the activation of ATP sensitive K^+^ channel, which facilitated membrane repolarization. By introducing the ATP sensitive K^+^ channel current into the Rasmusson’s cardiac model [17], we found the decay τ remained unchanged when ATP was greater than 0.02 mM inside the cell, while τ dramatically decreased upon ATP depletion (Figure 2f). The simulation result was in line with the fact that hypoxic conditions, which led to energy depletion, shortened the repolarization period of cardiac action potentials (Figure 2c). Moreover, the ATP sensitive K^+^ channel is blocked in high ATP level [22] supported the simulation result that decay τ remained relative constant when ATP was more than 1 mM. The shortening of action potential inhibited contractility [23] which could reduce the energy consumption of cardiomyocytes, potentially protecting the cell under energy depletion [24,25].

Optogenetic tools, ChR2(H134R) and OptoXR-β2AR, activated cardiomyocytes through different mechanisms. ChR2(H134R) induced membrane depolarization via cation influx, mainly Na^+^ and Ca^2+^. In contrast, OptoXR-β2AR increased intracellular cAMP and activated cAMP-PKA pathway [18]. cAMP affected the activity of HCN channels, which acted as pacemakers in heart [26]. PKA phosphorylates calcium channels to increase calcium influx [27], phosphorylates phospholamban to increase calcium reuptake into the sarcoplasmic reticulum [28], and phosphorylates troponin I and myosin binding protein C to increase calcium sensitivity and contractility [29]. The combined effects of these phosphorylation events are accelerated relaxation and contraction of cardiomyocytes, which increases the heart rate and pumping capacity.

Inhibitory rhodopsins, such as GR and ACR1, had distinct effects on cardiomyocytes. While they both inhibit cardiac beating, GR induced hyperpolarization whereas ACR1 did not. GR functions as an outward proton pump that increased cytosolic pH upon light illumination. Previous studies have reported that mitochondrial permeability transition pore (MPTP) opens when pH is higher than 7.3-7.5 and closes when pH is below 7[30]. Opening of MPTP causes calcium overload and cell death [31,32]. Prolonged light stimulation in GR-expressed cells alkalized the cytosol and might activate MPTP which led to calcium overload. Additionally, cytosolic alkalization can also inhibit the function of the calcium ATPase (SERCA) which is responsible for calcium reuptake to the sarco/endoplasmic reticulum, leading to an increase in cytosolic calcium levels [33]. This could explain the result that GR-expressing cardiomyocytes remained a sustained contracting state upon light illumination (Figure S1d). Previous reports showed that short low-intensity light pulses triggered action potential in ACR1 expressed cardiomyocytes, while sustained light cause depolarization and action potential inhibition [34]. Cardiomyocyte inhibition by sustained illumination polarized cells toward the reversal potential for Cl^-^, which is -40 mV∼ -33 mV[35], thus preventing cardiomyocytes repolarization [34].

Luminopsins (LMOs) are fusion proteins of luciferase and opsin that allow bioluminescence to activate light-driven channels and pumps [36]. Bioluminescence optogenetics enable modulating multiple brain regions without implanted hardware [37]. Currently available LMOs rely on marine luciferases, specifically either Gaussia luciferase (Gluc) or Renilla luciferase (Rluc), that use CTZ as their substrate [37]. CTZ is a hydrophobic molecule that can be administered to the brain using a variety of routes, such as intraperitoneal, intravenous, intracortical, or intranasal injection [37]. As visible light does not penetrate animal skin, traditional optogenetics in the heart requires invasive surgery or the use of far-red rhodopsin, which is limited in availability and lacks an inhibitory version so far. Bioluminescence optogenetics represents an alternative for controlling cardiac cell activity, offering distinct advantages over traditional methods.

Although chemogenetics tools such as hM3Dq and hM4Di have been applied in manipulating neuronal cell activities, cardiac chemogenetics have not yet been reported. The application of chemogenetics in cardiomyocytes, using hM3Dq and hM4Di, demonstrated a potential method for accelerating or inhibiting beating rates for a relatively long timescale compared to optogenetics. Furthermore, the establishment of an *in vitro* co-culture system with sympathetic neurons allowed the investigation of interactions between cardiomyocytes and sympathetic neurons. The sympathetic nervous system plays a crucial role in regulating cardiac function by increasing beat rate, contractile force, and conduction velocity[38]. Previous studies have developed methods for human-induced pluripotent stem cell-derived sympathetic neurons (hiPSC-SNs) and cardiomyocytes (hiPSC-CMs) co-culture [39], as well as hiPSC-CMs and primary mouse embryonic SNs coculture[38]. In our co-culture system, the activation of sympathetic neurons by hM3Dq led to an increased beating of cardiomyocytes, indicating that cardiomyocytes were successfully innervated by sympathetic neurons *in vitro*. hM3Dq activates the Gq pathway, leading to stimulation of phospholipase C, which catalyzes the conversion of phosphatidylinositol 4,5-bisphosphate to IP3 and DAG. Previous reports suggested that applying Gq-coupled agonists or IP3 caused an increase in firing rate of the adult mice hearts [40]. Optogenetic Gq activation by melanopsin in spontaneously beating embryoid bodies (EBs) increased their beating rate [41]. The result that Gq activation by hM3Dq increased the beating of cardiomyocytes was in line with these previous reports.

On the other hand, the Gi pathway, which inhibits the adenylate cyclase, decreases the cAMP level and reduces L-type Ca^2+^ current. Optogenetic Gi activation by LWO leads to an inhibition of spontaneously beating EBs [42]. hM4Di activates Gi pathway and decreases the beating of cardiomyocytes. However, the inhibitory effect of hM4Di in cardiomyocytes was limited, as the cardiomyocytes recovered their beating in a relatively short period. Consistently, *in vivo* ECG measurements showed that while hM3Dq increased heart beating rates in mice, hM4Di had a limited effect. This suggested that there might be alternative pathways that regulate cardiomyocyte beating when heart beating is inhibited by Gi activation, especially considering that the beating of cardiomyocytes is vital for animal survival.

In conclusion, this work offered comprehensive approaches to understanding the physiology of cardiomyocytes and manipulating heart beating rates. These methods include STFT analysis to extract the frequency of beating cardiomyocytes, mathematical model to simulate cardiomyocytes’ electrophysiological alterations under energy deficiency, cardiac-neuronal coculture, optogenetics and chemogenetics applications *in vivo* and *in vitro*. This methodology provides a comprehensive toolbox for studying the physiology of cardiomyocytes and potentially contributes to the drug development for cardiovascular diseases.

## 4. Materials and Methods

### 4.1 Plasmid construction

The sequence of GCAMP7s, hM3Dq, hM4Di, hChR2(H134R), OptoXR-b2AR, ACR1 and Gluc were obtained from Addgene (plasmid IDs: 104487, 50476, 50475, 20940, 20948, 67795, 114101). To enable expression in cardiomyocytes, cTnT promotor (-1 ∼ -589) was amplified from mouse genome DNA. For expression in neuron, hSyn promotor was amplified from plasmid (Addgene plasmid ID 114101). For AAV virus packaging, genes with promotors were cloned into pAAV-MCS vectors. All the constructions are verified by sequencing (Genewiz).

### 4.2 Primary cardiomyocyte isolation and culture

Postnatal day 0 (P0) C57BL mice hearts were dissected and cut into small pieces, which were then washed with HBSS. To digest the tissues, trypsin (Sigma-Aldrich) was applied for 5 minutes, followed by collagenase II (Sigma-Aldrich) for an additional 30 minutes. The cells were dissociated further using fire-polished pipettes before being plated onto coverslips (Glasswarenfabrik Karl Hecht, Germany, 12 mm) coated with Matrixgel (Corning). Cardiomyocytes were cultured in plating medium(100 ml of plating medium: 89 ml Minimal Essential Medium (Invitrogen), 0.5 g glucose, 0.5 mM glutamine, 2 g NaHCO3, 10 mg bovine transferrin (Calbiochem), 2.5 mg insulin, 10% FBS) and fed twice a week. Within 24 hours, the cardiomyocytes began to beat autonomously. For transient transfection, the isolated cardiomyocytes were subjected to electroporation before plating onto coverslips. Electrophysiology recordings were performed on cardiomyocytes at day in vitro (DIV) 3-4. For AAV transduction, the virus was added to the medium at DIV2-3 and imaged at DIV7-8.

### 4.3 Electroporation

Electroporation was employed to for transient transfection. Isolated cardiomyocytes were washed with PBS, centrifuged at 400 g for 5 minutes, and then resuspended in 700 µl of the electroporation solution (20 mM HEPES, 135 mM KCl, 2 mM MgCl_2_, 0.5% Ficol 400, 1% DMSO at pH 7.6, supplemented with 0.2 mM ATP and 0.5 mM Glutathione). The cells were transferred to a BTX disposable cuvette (4 mm gap). Plasmid DNA (20-30 μg) was added to the cells and kept on ice. The parameters of electroporation were set as follows: 180 V, 10 ms pulse duration, 2 pulses, and a 1-second interval. Following electroporation, cells were quickly transferred to a 15 ml tube, supplemented with 400 µl of FBS, and then centrifuged at 400 rpm for 5 minutes. Finally, cells were resuspended in plating medium and cultured according to standard procedures.

### 4.4 Cardiomyocyte electrophysiology and calcium imaging

Whole-cell patch clamp recordings were performed using a customized opto-electro system that included an Axopatch 700B amplifier (Molecular Devices), a Digidata 1440A (Molecular Devices) digitizer, an Optoscan (Cairn Research Ltd., UK) monochromator and an imaging system (Olympus) at room temperature. A customized Micro-Manager was used to control the devices. Data was acquired at a sampling rate of 10 kHz. Micropipettes were pulled from filamented glass capillaries (Sutter Instrument, BF150-86-10) using a micropipette puller (Sutter Instrument, P1000) to produce a tip resistance of 5-8 MΩ. The micropipette was filled with intracellular buffer (potassium gluconate 120 mM, KCl 3 mM, HEPES 10 mM, NaCl 8 mM, CaCl_2_ 0.5 mM EGTA 5 mM, ATP-Mg 2 mM, GTP 0.3 mM, pH 7.2) and positioned using a micromanipulator (MP285, Sutter). Cardiomyocytes were loaded with Fluo-4-AM (Invitrogene) to a final concentration of 5 μM in Tyrode’s buffer (NaCl 145 mM, KCl 3 mM, HEPES 10 mM, glucose 10 mM, pH 7.4) for 15 min at 37. For calcium imaging, current-clamp mode was used to record membrane potential and cells were imaged by a fluorescence microscope (Olympus IX83, Japan) with a 40x water objective in Tyrode’s buffer. The fluorescence signal was excited with 480 nm with a bandwidth of 10 nm light generated by monochromator and emission light was collected with a 520-560 nm emission fitter. A customized protocol was established, which included a digital signal triggering the camera for image acquisition. Images are acquired throughout current-clamp recording.

### 4.5 Electrical stimulation

The cardiomyocytes, cultured on a coverslip, were placed in a chamber with a pair of parallel metal wires distributed on either side of the chamber. A customized electrical stimulation device (Arduino) was used to give 5 V voltage pulses on the two metal wires. This distance between two wires was 7 mm. Cardiomyocytes stained with Fluo-4-AM were imaged by a fluorescence microscope (Olympus IX83, Japan) in Tyrode’s buffer. Images are acquired throughout the electrical stimulation process.

### 4.6 Drug treatment and living cells imaging

Fluo-4-stained cells were observed under a fluorescence microscope (Olympus IX83, Japan) in Tyrode’s buffer. Cells were treated with 2DG (10 mM) to inhibit glycolysis. Oligomycin (100 ng/ml) or NaN_3_ (0.2 mg/ml) was used to inhibit oxidative phosphorylation, while CCCP (50 μM) was utilized to uncouple the mitochondrial potential of cardiomyocytes. To capture the voltage dynamics during energy depletion, cardiomyocytes were clamped using current-clamp mode, and their activities were recorded throughout the 2DG treatment process.

### 4.7 Virus preparation and transduction

For the transfection of cultured cells, we employed AAV DJ serotype virus. To generate this virus, gene expression plasmid, capsid (pAAV-DJ), and helper plasmids (pHelper) were co-transfected into 293t cells using the calcium phosphate precipitation method. Following a 48-hour incubation period, viruses were collected from cell pellets via four cycles of frozen-thaw methods. The titer of the virus was determined through real-time PCR analysis.

### 4.8 Optogenetic stimulation

Cells transfected with ChR2(H134R) were stimulated by blue light pulses (473 nm, 10 mW) with timescale ranging from 20-500 ms through a 40x objective (Olympus) in Tyrode’s buffer. Patch clamping was used to record the membrane potential during light stimulation, and bright field images were acquired simultaneously. For light activation with Opto-XR β2AR, cardiomyocytes were loaded with Fura-2 (Invitrogene) to a final concentration of 5 μM with 0.04% F127 (Sigma-Aldrich) in Tyrode’s buffer for 15 min at 37. Fura-2 was excited by 340 nm and 380 nm, and emission was collected at 520 ± 20 nm. Cells are illuminated by green light (532 nm, 10 mW). The [Ca^2+^] was determined by calculating the 340 nm/380 nm ratio of the fluorescence images. For light inhibition with GR or ACR1, cardiomyocytes were illuminated with green light (532 nm, 10 mW) and the bright field images were acquired during light stimulation. Membrane potential of cardiomyocyte was recorded via patch clamping during light stimulation.

### 4.9 Auto-illuminance stimulation

Cardiomyocytes transfected with Gluc-ACR1 were stimulated by the addition of CTZ (MCE) to a final concentration of 100 mM. To confirm illuminance, cells cultured on a coverslip were placed in a chamber in Tyrode’s buffer and imaged using a cooled-CCD camera after CTZ application. Bright field images were acquired during the process of CTZ addition to observe auto-illuminance induced changes in the cardiomyocytes’ beating.

### 4.10 Video-based cardiomyocytes beating analysis

According to the Nyquist sampling rule, the sampling frequency for imaging should be more than 2-fold that of the cardiomyocytes’ beating frequency *in vitro*. Bright field images were acquired using an EMCCD camera (Photometric Evolve) at a sampling rate of 8 Hz. The acquired video was analyzed using ImageJ (NIH). First, the video was converted to 8-bit and a line was drawn on the video to cover the movement of cells. A Kymograph was then generated using the Multiple Kymograph plugin. By drawing a straight line across the time-axis of the kymograph, a plot of grayscale-value versus time was generated using the “Plot Profile” command. The grayscale-value data was saved and used for spectrogram analysis. A customized Python script was utilized to generate the spectrogram. The scripts were provided in the supplementary.

### 4.11 Chemogenetics stimulation

Cardiomyocytes transfected with hM3Dq or hM4Di and GCAMP7s were imaged by a fluorescence microscope (Olympus IX83, Japan) in Tyrode’s buffer. CNO (MCE) was added to the solution to a final concentration of 2 μM.

### 4.12 Cardiomyocyte and sympathetic neuron coculture

Sympathetic neurons were isolated from the spinal cord of postnatal day 0 (P0) C57BL mice. Trypsin was used to break down the cells for five minutes, after which single cells were produced through further breakdown using fire-polished pipettes. The cells were then harvested by centrifugation, mixed with isolated cardiomyocytes, and plated onto matrix gel-coated coverslips. The cells were cultured in plating medium and fed twice a week. Virus transduction was performed at DIV3, and the cells were imaged at DIV7-8.

### 4.13 Virus injection

Adult mice weighing 24-26 g were anesthetized with isoflurane and injected with AAV virus to the angular vein using a 29G 1 ml insulin syringe (BD). Following injection, mice were further raised for 4-8 weeks allowing for virus infection and expression.

### 4.14 ECG recording and CNO injection

Mice injected with virus were anesthetized with 3% pentobarbital sodium (intraperitoneal injection, 30 mg/kg). ECG was recorded by a 4-lead ECG system (iWorx, BIO4). Electrodes were plugged in the four limbs of the mouse according to the manufactures’ instruction. CNO was injected intraperitoneally with a dose of 2 mg/kg. ECG was recorded for 1 hour after injection.

### 4.15 Action potential modeling

The original Rasmusson model [17] was modified to include the current of the membrane ATP sensitive potassium channel (I_KATP_), which was not present in the original model. The equations and parameters for I_KATP_ were obtained from the Luo Rudy dynamic model [43]. The resulting cardiomyocyte action potential model was formatted in ode form and can be simulated using the XPPAUT program. To investigate the effect of ATP concentration on the decay τ of action potential, the intracellular ATP concentration parameter (ATPi) was adjusted from 3 mM to 3 μM. The curve from the peak of the action potential to the 20% amplitude value was fitted using a single exponential decay equation (equation 1).

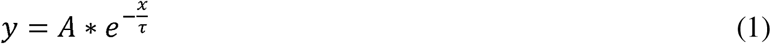

All the equations and parameters were listed in the supplementary.

### 4.16 Data and statistical analysis

All electrophysiology data were analyzed using pClampfit (Molecular Devices) in combination with customized Perl scripts. For patch clamping and calcium imaging analysis, the images obtained were analyzed using ImageJ, with fluorescence curves being aligned to calcium curves. Kymographs were generated through the ImageJ MultipleKymograph plugin. Statistical analysis was performed with unpaired two-tailed Student’s t tests being used for comparison of two samples.

## Supporting information

Supplementary Information_cardiomyocytes

## Author Contributions

Conceptualization, R.Z.Y.; methodology, R.Z.Y.; software, R.Z.Y.; validation, R.Z.Y., D.H.L. and S.M.L.; formal analysis, R.Z.Y.; investigation, R.Z.Y.; resources, J.S.K., P.P.L. and S.A.L.; data curation, R.Z.Y.; writing—original draft preparation, R.Z.Y.; writing—review and editing, R.Z.Y.; visualization, R.Z.Y.; supervision, J.S.K.; project administration, J.S.K.; funding acquisition, R.Z.Y, J.S.K., P.P.L. and S.A.L.

## Animal Ethics

The study was conducted according to the guidelines of the Institutional Animal Care and Use Committee of the Zhengzhou University under project ID 2024-KY-0399-001, April 2024.

## Data Availability Statement

Data are available upon request from the authors.

## Acknowledgments

The author gratefully acknowledges the support provided by the First Affiliated Hospital of Zhengzhou University, Zhengzhou, Henan province, PRC.

