## Supplementary Information_cardiomyocytes for "Methodologies for Manipulating Cardiomyocyte Physiology: *In Vitro* and *In Vivo* Perspectives"

Supplementary Figures

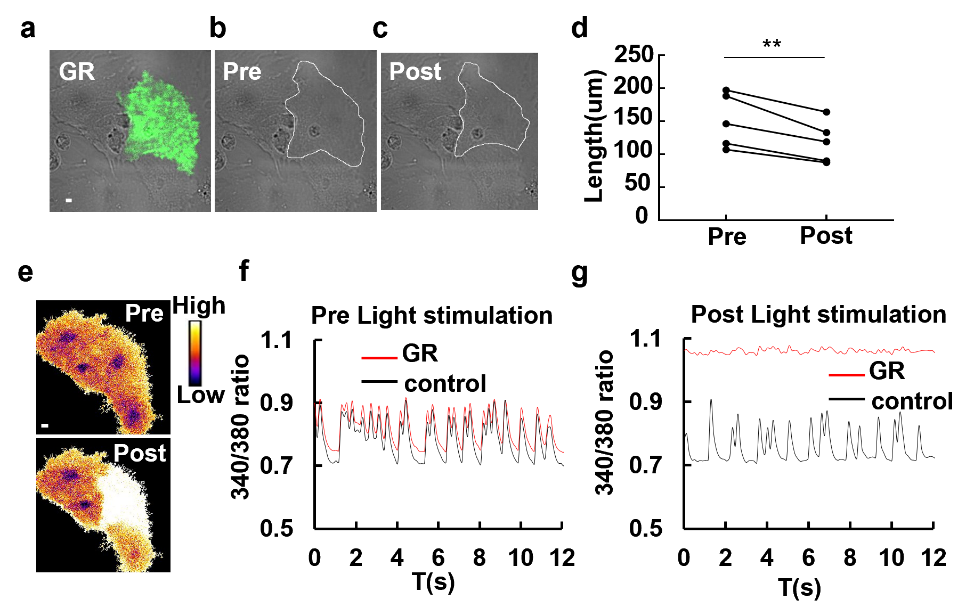

Figure S1. Prolonged light exposure leads to calcium overload in cardiomyocytes expressing GR. (a) Image of cardiomyocytes transfected with GR. Scale bar, 5 μm. (b) Bright filed image of cardiomyocytes before light stimulation. The outline of the GR-transfected cell was marked with white line. (c) Bright filed image of cardiomyocytes after light stimulation. The outline of the GR-transfected cell was marked with white line. (d) Diameter of cardiomyocytes transfected with GR before and after light stimulation. There was a significant decrease in cell length upon light stimulation (*Paired t-test*, *p* = 0.0060). (e) Calcium images of cardiomyocytes transfected with GR before (top) and after (bottom) light stimulation. Calcium concentration was indicated by Fura-2 ratio represented in pseudo-color. This image is the same filed in (a). Scale bar, 5 μm. (f) Quantification of intracellular Ca^2+^ concentration of cardiomyocytes transfected with GR (red) and control (black) before light stimulation. (g) Quantification of intracellular Ca^2+^ concentration of cardiomyocytes transfected with GR (red) and control (black) after light stimulation.

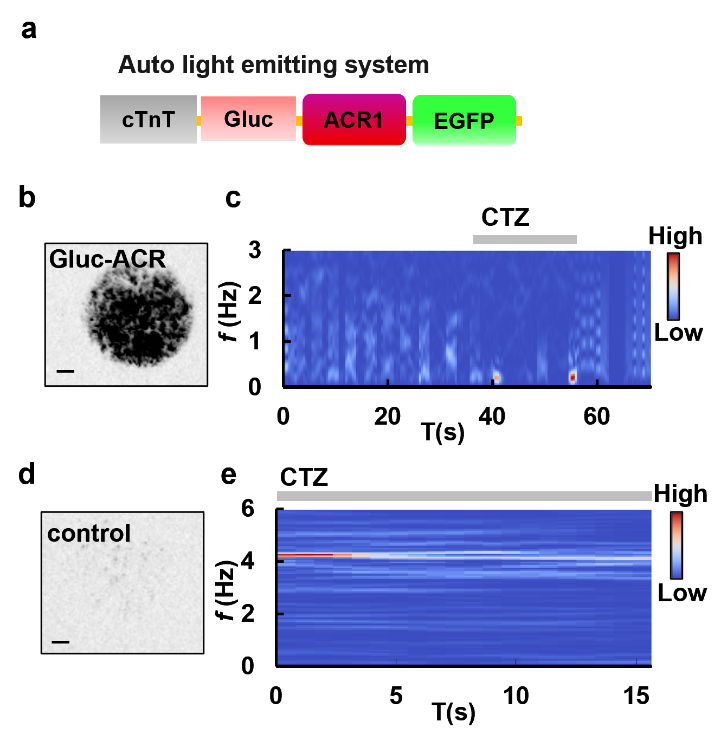

Figure S2. Bioluminescence inhibition of beatings of cardiomyocytes. (a) The construction of bioluminescence activated cardiomyocytes inhibition system. cTnT promoter was used to drive specific expressions in cardiomyocytes. The luciferase Gluc was fused to the N-terminus of ACR1 and EGFP was fused to the C-terminus of ACR1. (b) Chemiluminescence imaging of cardiomyocytes cultured on a coverslip. CTZ was added to induce chemiluminescence. Scale bar, 2 mm. The circular outline indicates the shape of the coverslip. (c) Spectrogram of cardiomyocytes expressing Gluc-ACR1 before and after addition of CTZ based on STFT analysis of bright field video. Cardiomyocytes’ beating was inhibited after the addition of CTZ and resumed when CTZ was washed away. (d) Chemiluminescence imaging of control cardiomyocytes upon CTZ addition. No positive signals can be detected in cardiomyocytes without Gluc-ACR1-EGFP expression. Scale bar, 2 mm. (e) Spectrogram of control cardiomyocytes before and after addition of CTZ based on bright field video analysis.

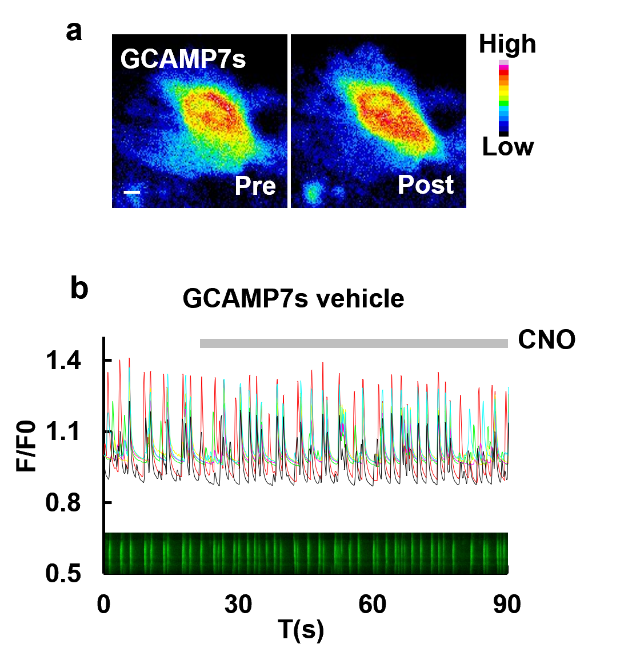

Figure S3. CNO itself has no effects on the beatings of cardiomyocytes. (a) Calcium imaging of cardiomyocytes transfected with GCAMP7s before (left) and after (right) CNO addition. The image was presented as pseudo-color. Scale bar, 5 μm. (b) Quantification of intracellular Ca^2+^ concentration of cardiomyocytes upon CNO addition. The gray bar indicates the CNO application. The inset indicates the kymograph of intracellular Ca^2+^.
Supplementary Movies

**Supplementary Movie 1.** Calcium dynamics in cardiomyocytes. Scale bar, 5 μm. Cardiomyocytes were stained with Fluo-4. Images were represented in pseudo-color.

**Supplementary Movie 2.** Calcium flooding in cardiomyocytes. Scale bar, 5 μm. Cardiomyocytes were stained with Fluo-4. Images were represented in pseudo-color.

**Supplementary Movie 3.** Calcium sparks in cardiomyocytes. Scale bar, 5 μm. Cardiomyocytes were stained with Fluo-4. Images were represented in pseudo-color.

**Supplementary Movie 4.** Calcium dynamics in cardiomyocytes upon electrical stimulation. Scale bar, 5 μm. Cardiomyocytes were stained with Fluo-4.

**Supplementary Movie 5.** Calcium dynamics in cardiomyocytes upon 2DG treatment. Scale bar, 5 μm. Cardiomyocytes were stained with Fluo-4.

**Supplementary Movie 6.** Calcium dynamics in cardiomyocytes upon oligomycin treatment. Scale bar, 5 μm. Cardiomyocytes were stained with Fluo-4.

**Supplementary Movie 7.** Calcium dynamics in cardiomyocytes upon NaN_3_ treatment. Scale bar, 5 μm. Cardiomyocytes were stained with Fluo-4.

**Supplementary Movie 8.** Calcium dynamics in cardiomyocytes upon CCCP treatment. Scale bar, 5 μm. Cardiomyocytes were stained with Fluo-4.

**Supplementary Movie 9.** Bright field video in cardiomyocytes transfected with ChR2(H134R)-EGFP upon light stimulation. Scale bar, 5 μm.

**Supplementary Movie 10.** Calcium dynamics in cardiomyocytes transfected with OptoXR-β2AR-EGFP before and after light stimulation. Scale bar, 5 μm. Cardiomyocytes were stained with Fura-2. Images were represented in pseudo-color.

**Supplementary Movie 11.** Bright field video in cardiomyocytes transfected with GR-EGFP upon light stimulation. Scale bar, 5 μm.

**Supplementary Movie 12.** Calcium dynamics in cardiomyocytes transfected with GR-EGFP before and after light stimulation. Scale bar, 5 μm. Cardiomyocytes were stained with Fura-2. The cell expressing GR was indicated by green arrow. Images were represented in pseudo-color.

**Supplementary Movie 13.** Bright field video in cardiomyocytes transfected with ACR1-EGFP upon light stimulation. Scale bar, 5 μm.

**Supplementary Movie 14.** Calcium dynamics in cardiomyocytes transfected with cTnT-hM3D-mcherry and cTnT-GCAMP7s upon CNO treatment. Scale bar, 5 μm.

**Supplementary Movie 15.** Calcium dynamics in cardiomyocytes transfected with cTnT-hM4D-mcherry and cTnT-GCAMP7s upon CNO treatment. Scale bar, 5 μm.

**Supplementary Movie 16.** Calcium dynamics in cardiomyocytes with cTnT-GCAMP7s upon CNO treatment. Scale bar, 5 μm.

**Supplementary Movie 17.** Calcium dynamics in co-culture of cardiomyocytes and neurons transfected with hSyn-hM3D-mcherry and cTnT-GCAMP7s upon CNO treatment. Scale bar, 5 μm.

**Supplementary Movie 18.** Calcium dynamics in cardiomyocytes without neurons transfected with cTnT-GCAMP7s and hSyn-hM3D-mcherry upon CNO treatment. Scale bar, 5 μm.

Supplementary Text

The following equations (1-118) are used to build the model for cardiomyocytes.

$$\begin{aligned} E_{K}=\frac{RT}{F}\ln\frac{\left[ K_{o} \right]}{\left[ K_{i} \right]}\#\left( \right) \end{aligned}$$

$$C_{m}\frac{dV}{dt}=-I_{CaL}+I_{pCa}+I_{NaCa}+I_{Cab}+I_{Na}+I_{Nab}+I_{NaK}+I_{Kto,f}+I_{Kto,s}+I_{K1}+I_{Ks}+$$

$$\begin{aligned} I_{Kur}+I_{Kss}+I_{Kr}+I_{KATP}+I_{Cl,Ca}-r_{stat}\#\left( \right) \end{aligned}$$

$$\begin{aligned} \frac{d\left[ K_{i} \right]}{dt}=-I_{Kto,f}+I_{Kto,s}+I_{K1}+I_{Ks}+I_{Kss}+I_{Kur}+I_{Kr}-2I_{NaK}\frac{A_{cap}C_{m}}{V_{myo}F}\#\left( \right) \end{aligned}$$

$$\begin{aligned} I_{Kto,f}=G_{Kto,f}a_{to,f}^{3}i_{to,f}\left( V-E_{K} \right)\#\left( \right) \end{aligned}$$

$$\begin{aligned} \alpha_{a}=0.18064 e^{0.03577\left( V+30.0 \right)}\#\left( \right) \end{aligned}$$

$$\begin{aligned} \beta_{a}=0.3956 e^{-0.06237\left( V+30.0 \right)}\#\left( \right) \end{aligned}$$

$$\begin{aligned} \alpha_{i1}=\frac{0.000152 e^{-\left( V+13.5 \right)/7}}{0.0067083 e^{-\left( V+33.5 \right)/7}+1}\#\left( \right) \end{aligned}$$

$$\begin{aligned} \beta_{i1}=\frac{0.00095 e^{\left( V+33.5 \right)/7}}{0.051335 e^{\left( V+33.5 \right)/7}+1}\#\left( \right) \end{aligned}$$

$$\begin{aligned} \frac{da_{to,f}}{dt}=\alpha_{a}\left( 1-a_{to,f} \right)-\beta_{a}a_{to,f}\#\left( \right) \end{aligned}$$

$$\begin{aligned} \frac{di_{to,f}}{dt}=\alpha_{i1}\left( 1-i_{to,f} \right)-\beta_{i1}i_{to,f}\#\left( \right) \end{aligned}$$

$$\begin{aligned} I_{Kto,s}=G_{Kto,s}a_{to,s}i_{to,s}\left( V-E_{K} \right)\#\left( \right) \end{aligned}$$

$$\begin{aligned} \frac{da_{to,s}}{dt}=\frac{a_{ss}-a_{to_{,}s}}{\tau_{ta,s}}\#\left( \right) \end{aligned}$$

$$\begin{aligned} \frac{di_{to,s}}{dt}=\frac{i_{ss}-i_{to,s}}{\tau_{ti,s}}\#\left( \right) \end{aligned}$$

$$\begin{aligned} a_{ss}=\frac{1}{e^{-\left( V+22.5 \right)/7.7}+1}\#\left( \right) \end{aligned}$$

$$\begin{aligned} i_{ss}=\frac{1}{e^{-\left( V+45.2 \right)/5.7}+1}\#\left( \right) \end{aligned}$$

$$\begin{aligned} \tau_{ta,s}=2.058+0.493 e^{-0.0629V}\#\left( \right) \end{aligned}$$

$$\begin{aligned} \tau_{ti,s}=270.0+\frac{1050.0}{1+e^{\left( V+45.2 \right)/5.7}}\#\left( \right) \end{aligned}$$

$$\begin{aligned} I_{K1}=0.2938\left( \frac{\left[ K_{o}^{+} \right]}{\left[ K_{o}^{+} \right]+210.0} \right) \left[ \frac{V-E_{K}}{1+e^{0.0896\left( V-E_{K} \right)}} \right]\#\left( \right) \end{aligned}$$

$$\begin{aligned} I_{Ks}=G_{Ks}n_{Ks}^{2}\left( V-E_{K} \right)\#\left( \right) \end{aligned}$$

$$\begin{aligned} \frac{dn_{Ks}}{dt}=\alpha_{n}\left( 1-n_{Ks} \right)-\beta_{n}n_{Ks}\#\left( \right) \end{aligned}$$

$$\begin{aligned} \alpha_{n}=4.81333\times{10}^{-6}\left( 26.5+V \right)\left[ 1.0-e^{-0.128\left( V+26.5 \right)} \right]\#\left( \right) \end{aligned}$$

$$\begin{aligned} \beta_{n}=9.53333\times{10}^{-5}e^{-0.038\left( V+26.5 \right)}\#\left( \right) \end{aligned}$$

$$\begin{aligned} I_{Kur}=G_{Kur}a_{ur}i_{ur}\left( V-E_{K} \right)\#\left( \right) \end{aligned}$$

$$\begin{aligned} \frac{da_{ur}}{dt}=\frac{a_{ss}-a_{ur}}{\tau_{aur}}\#\left( \right) \end{aligned}$$

$$\begin{aligned} \frac{di_{ur}}{dt}=\frac{i_{ss}-i_{ur}}{\tau_{iur}}\#\left( \right) \end{aligned}$$

$$\begin{aligned} \tau_{aur}=2.058+0.493 e^{-0.0629V}\#\left( \right) \end{aligned}$$

$$\begin{aligned} \tau_{iur}=1200.0-\frac{170.0}{1.0+e^{\left( V+45.2 \right)/5.7}}\#\left( \right) \end{aligned}$$

$$\begin{aligned} I_{Kss}=G_{Kss}a_{Kss}i_{Kss}\left( V-E_{K} \right)\#\left( \right) \end{aligned}$$

$$\begin{aligned} \frac{da_{Kss}}{dt}=\frac{a_{ss}-a_{Kss}}{\tau_{Kss}}\#\left( \right) \end{aligned}$$

$$\begin{aligned} \frac{di_{Kss}}{dt}=0\#\left( \right) \end{aligned}$$

$$\begin{aligned} \tau_{Kss}=13.17+39.3e^{-0.0862V}\#\left( \right) \end{aligned}$$

$$\begin{aligned} I_{Kr}=G_{Kr}O_{K}\left[ V-\frac{RT}{F} \ln\left( \frac{0.98\left[ K^{+} \right]_{o}+0.02\left[ Na^{+} \right]_{o}}{0.98\left[ K^{+} \right]_{i}+0.02\left[ Na^{+} \right]_{i}} \right) \right]\#\left( \right) \end{aligned}$$

$$\begin{aligned} C_{K0}=1-\left( C_{K1}+C_{K2}+O_{K}+I_{K} \right)\#\left( \right) \end{aligned}$$

$$\begin{aligned} \frac{dC_{K1}}{dt}=-\left( \beta_{a0}C_{K1}+k_{f}C_{K1} \right)+\alpha_{a0}C_{K0}+k_{b}C_{K2}\#\left( \right) \end{aligned}$$

$$\begin{aligned} \frac{dC_{K2}}{dt}=-\left( k_{b}C_{K2}+\alpha_{a1}C_{K2} \right)+\beta_{a1}O_{K}+k_{f}C_{K1}\#\left( \right) \end{aligned}$$

$$\begin{aligned} \frac{dO_{K}}{dt}=-\left( \beta_{a1}O_{K}+\alpha_{i}O_{K} \right)+\alpha_{a1}C_{K2}+\beta_{i}I_{K}\#\left( \right) \end{aligned}$$

$$\begin{aligned} \frac{dI_{K}}{dt}=\alpha_{i}O_{K}-\beta_{i}I_{K}\#\left( \right) \end{aligned}$$

$$\begin{aligned} \alpha_{a0}=0.022348e^{0.01176V}\#\left( \right) \end{aligned}$$

$$\begin{aligned} \alpha_{a1}=0.013733e^{0.038198V}\#\left( \right) \end{aligned}$$

$$\begin{aligned} \beta_{a0}=0.047002e^{-0.0631V}\#\left( \right) \end{aligned}$$

$$\begin{aligned} \beta_{a1}=6.89\times{10}^{-5}e^{-0.04178V}\#\left( \right) \end{aligned}$$

$$\begin{aligned} \alpha_{i}=0.090821e^{0.023391\left( V+5.0 \right)}\#\left( \right) \end{aligned}$$

$$\begin{aligned} \beta_{i}=0.006497e^{-0.03268\left( V+5.0 \right)}\#\left( \right) \end{aligned}$$

$$\begin{aligned} I_{NaK}=I_{NaK}^{max}f_{NaK}\left( \frac{\left[ K \right]_{o}}{\left[ K \right]_{o}+K_{m,Ko}} \right)\cdot\{\frac{1}{1+\left( K_{m,Nai}/\left[ Na^{+} \right]_{i} \right)^{3/2}}\}\#\left( \right) \end{aligned}$$

$$\begin{aligned} f_{NaK}=\frac{1}{1+0.1245 e^{-0.1VF/RT}+0.0365 \sigma e^{-0.1VF/RT}}\#\left( \right) \end{aligned}$$

$$\begin{aligned} \sigma=\frac{1}{7}\left( e^{\left[ Na^{+} \right]_{o}/67300}-1 \right)\#\left( \right) \end{aligned}$$

$$\begin{aligned} E_{Na}=\frac{RT}{F} \ln\frac{0.9\left[ Na^{+} \right]_{o}+0.1\left[ K^{+} \right]_{o}}{0.9\left[ Na^{+} \right]_{i}+0.1\left[ K^{+} \right]_{i}}\#\left( \right) \end{aligned}$$

$$\begin{aligned} \frac{d\left[ Na^{+} \right]_{i}}{dt}=-\frac{A_{cap}C_{m}}{V_{myo}F}\left( I_{Na}+I_{Nab}+3I_{NaK}+3I_{NaCa} \right)\#\left( \right) \end{aligned}$$

$$\begin{aligned} I_{Na}=G_{Na}O_{Na}\left( V-E_{Na} \right)\#\left( \right) \end{aligned}$$

$$\begin{aligned} E_{Na}=\frac{RT}{F}\ln\left( \frac{0.9\left[ Na^{+} \right]_{o}+0.1\left[ K^{+} \right]_{o}}{0.9\left[ Na^{+} \right]_{i}+0.1\left[ K^{+} \right]_{i}} \right)\#\left( \right) \end{aligned}$$

$$\begin{aligned} C_{Na3}=1.0-\left( O_{Na}+C_{Na1}+C_{Na2}+IF_{Na}+I1_{Na}+I2_{Na}+IC_{Na2}+IC_{Na3} \right)\#\left( \right) \end{aligned}$$

$$\begin{aligned} \frac{dC_{Na2}}{dt}=-\left( \beta_{Na11}C_{Na2}+\alpha_{Na12}C_{Na2}+\beta_{Na3}C_{Na2} \right) +\alpha_{Na11}C_{Na3}+\beta_{Na12}C_{Na1}+\alpha_{Na3}IC_{Na2}\#\left( \right) \end{aligned}$$

$$\begin{aligned} \frac{dC_{Na1}}{dt}=-\left( \beta_{Na12}C_{Na1}+\alpha_{Na13}C_{Na1}+\beta_{Na3}C_{Na1} \right) +\alpha_{Na12}C_{Na2}+\beta_{Na13}O_{Na}+\alpha_{Na3}IF_{Na}\#\left( \right) \end{aligned}$$

$$\begin{aligned} \frac{dO_{Na}}{dt}=-\left( \beta_{Na13}O_{Na}+\alpha_{Na2}O_{Na} \right)+\alpha_{Na13}C_{Na1}+\beta_{Na2}IF_{Na}\#\left( \right) \end{aligned}$$

$$\frac{dIF_{Na}}{dt}=-\left( \beta_{Na2}IF_{Na}+\alpha_{Na3}IF_{Na}+\alpha_{Na4}IF_{Na}+\beta_{Na12}IF_{Na} \right)+$$

$$\begin{aligned} \alpha_{Na2}O_{Na}+\beta_{Na3}C_{Na1}+\beta_{Na4}I1_{Na}+\alpha_{Na12}IC_{Na2}\#\left( \right) \end{aligned}$$

$$\begin{aligned} \frac{dI1_{Na}}{dt}=-\left( \beta_{Na4}I1_{N}a+\alpha_{Na5}I1_{Na} \right)+\alpha_{Na4}IF_{Na}+\beta_{Na5}I2_{Na}\#\left( \right) \end{aligned}$$

$$\begin{aligned} \frac{dI2_{Na}}{dt}=\alpha_{Na5}I1_{Na}-\beta_{Na5}I2_{Na}\#\left( \right) \end{aligned}$$

$$\frac{dIC_{Na2}}{dt}=-\left( \beta_{Na11}IC_{Na2}+\alpha_{Na12}IC_{Na2}+\alpha_{Na3}IC_{Na2} \right) +$$

$$\begin{aligned} \alpha_{Na11}IC_{Na3}+\beta_{Na12}IF_{Na}+\beta_{Na13}IC_{Na2}\#\left( \right) \end{aligned}$$

$$\frac{dIC_{Na3}}{dt}=-\left( \alpha_{Na11}IC_{Na3}+\alpha_{Na3}IC_{Na3} \right) +$$

$$\begin{aligned} \beta_{Na11}IC_{Na2}+\beta_{Na3}C_{Na3}\#\left( \right) \end{aligned}$$

$$\begin{aligned} \alpha_{Na11}=\frac{3.802}{0.1027e^{-\left( V+2.5 \right)/17}+0.2e^{-\left( V+2.5 \right)/150}}\#\left( \right) \end{aligned}$$

$$\begin{aligned} \alpha_{Na12}=\frac{3.802}{0.1027e^{-\left( V+2.5 \right)/15}+0.23e^{-\left( V+2.5 \right)/150}}\#\left( \right) \end{aligned}$$

$$\begin{aligned} \alpha_{Na13}=\frac{3.802}{0.1027e^{-\left( V+2.5 \right)/12}+0.25e^{-\left( V+2.5 \right)/150}}\#\left( \right) \end{aligned}$$

$$\begin{aligned} \beta_{Na11}=0.1917 e^{-\left( V+2.5 \right)/20.3}\#\left( \right) \end{aligned}$$

$$\begin{aligned} \beta_{Na12}=0.2 e^{-\left( V-2.5 \right)/20.3}\#\left( \right) \end{aligned}$$

$$\begin{aligned} \beta_{Na13}=0.22 e^{-\left( V-7.5 \right)/20.3}\#\left( \right) \end{aligned}$$

$$\begin{aligned} \alpha_{Na3}=7.0\times{10}^{-7}e^{-\left( V+7 \right)/7.7}\#\left( \right) \end{aligned}$$

$$\begin{aligned} \beta_{Na3}=0.0084+2.0\times{10}^{-5}\left( V+7.0 \right)\#\left( \right) \end{aligned}$$

$$\begin{aligned} \alpha_{Na2}=\frac{1.0}{0.393956+0.188495 e^{\left( V+7.0 \right)/16.6}}\#\left( \right) \end{aligned}$$

$$\begin{aligned} \beta_{Na2}=\alpha_{Na13} \alpha_{Na2} \alpha_{Na3}/\left( \beta_{Na13} \beta_{Na3} \right)\#\left( \right) \end{aligned}$$

$$\begin{aligned} \alpha_{Na4}=0.001\alpha_{Na2}\#\left( \right) \end{aligned}$$

$$\begin{aligned} \beta_{Na4}=\alpha_{Na3}\#\left( \right) \end{aligned}$$

$$\begin{aligned} \alpha_{Na5}=\alpha_{Na2}/95000\#\left( \right) \end{aligned}$$

$$\begin{aligned} \beta_{Na5}=0.02\alpha_{Na3}\#\left( \right) \end{aligned}$$

$$\begin{aligned} I_{Nab}=G_{Nab}\left( V-E_{Na} \right)\#\left( \right) \end{aligned}$$

$$\begin{aligned} E_{CaN}=\frac{RT}{2F}\ln\frac{\left[ Ca^{2+} \right]_{o}}{\left[ Ca^{2+} \right]_{i}}\#\left( \right) \end{aligned}$$

$$\begin{aligned} I_{CaL}=G_{CaL}O\left( V-E_{CaL} \right)\#\left( \right) \end{aligned}$$

$$\frac{d\left[ Ca^{2+} \right]_{i}}{dt}=B_{i}\{J_{leak}+J_{xfer}-J_{up}-J_{trpn}-$$

$$\begin{aligned} \left( I_{Cab}+I_{pCa}-2I_{NaCa} \right)\frac{A_{cap}C_{m}}{2V_{myo}F}\}\#\left( \right) \end{aligned}$$

$$\begin{aligned} B_{i}=\{1+\frac{\left[ CMDN \right]_{tot}K_{m}^{CMDN}}{K_{m}^{CMDN}+\left[ Ca^{2+} \right]_{i}^{2}}{\}}^{-1}\#\left( \right) \end{aligned}$$

$$\begin{aligned} B_{ss}=\{1+\frac{\left[ CMDN \right]_{tot}K_{m}^{CMDN}}{K_{m}^{CMDN}+\left[ Ca^{2+} \right]_{ss}^{2}}{\}}^{-1}\#\left( \right) \end{aligned}$$

$$\begin{aligned} B_{JSR}=\{1+\frac{\left[ CSQN \right]_{tot}K_{m}^{CSQN}}{K_{m}^{CSQN}+\left[ Ca^{2+} \right]_{JSR}^{2}}{\}}^{-1}\#\left( \right) \end{aligned}$$

$$\begin{aligned} \frac{d\left[ Ca^{2+} \right]_{ss}}{dt}=B_{ss}\left( J_{rel}\frac{V_{JSR}}{V_{ss}}-J_{xfer}\frac{V_{myo}}{V_{ss}}-I_{CaL}\frac{A_{cap}C_{m}}{2V_{ss}F} \right)\#\left( \right) \end{aligned}$$

$$\begin{aligned} \frac{d\left[ Ca^{2+} \right]_{JSR}}{dt}=B_{JSR}\left( J_{tr}-J_{rel} \right)\#\left( \right) \end{aligned}$$

$$\begin{aligned} \frac{d\left[ Ca^{2+} \right]_{NSR}}{dt}=\left( J_{up}-J_{leak} \right)\frac{V_{myo}}{V_{NSR}}-J_{tr}\frac{V_{JSR}}{V_{NSR}}\#\left( \right) \end{aligned}$$

$$\begin{aligned} J_{rel}=\nu_{1}\left( O_{1}+O_{2} \right)\left( \left[ Ca^{2+} \right]_{JSR}-\left[ Ca^{2+} \right]_{ss} \right)P_{RyR}\#\left( \right) \end{aligned}$$

$$\begin{aligned} J_{leak}=\nu_{2}\left( \left[ Ca^{2+} \right]_{NSR}-\left[ Ca^{2+} \right]_{i} \right)\#\left( \right) \end{aligned}$$

$$\begin{aligned} J_{xfer}=\frac{\left[ Ca^{2+} \right]_{ss}-\left[ Ca^{2+} \right]_{i}}{\tau_{xfer}}\#\left( \right) \end{aligned}$$

$$\begin{aligned} J_{up}=\nu_{3}\frac{\left[ Ca^{2+} \right]_{i}^{2}}{K_{m,up}^{2}+\left[ Ca^{2+} \right]_{i}^{2}}\#\left( \right) \end{aligned}$$

$$\begin{aligned} J_{tr}=\frac{\left[ Ca^{2+} \right]_{NSR}-\left[ Ca^{2+} \right]_{JSR}}{\tau_{tr}}\#\left( \right) \end{aligned}$$

$$J_{trpn}=-\left( k_{htrpn}^{-}\left[ HTRPNCa \right]+k_{ltrpn}^{-}\left[ LTRPNCa \right] \right) +$$

$$k_{htrpn}^{+}\left[ Ca^{2+} \right]_{i}\left( \left[ HTRPN \right]_{tot}-\left[ HTRPNCa \right] \right)$$

$$\begin{aligned} +k_{ltrpn}^{+}\left[ Ca^{2+} \right]_{i}\left( \left[ LTRPN \right]_{tot}-\left[ LTRPNCa \right] \right)\#\left( \right) \end{aligned}$$

$$\begin{aligned} \frac{dP_{RyR}}{dt}=-0.04P_{RyR}-0.1\frac{I_{CaL}}{I_{CaL,max}}e^{-\frac{\left( V-5.0 \right)^{2}}{648}}\#\left( \right) \end{aligned}$$

$$\begin{aligned} \frac{d\left[ LTRPNCa \right]}{dt}=k_{ltrpn}^{+}\left[ Ca^{2+} \right]_{i}\left( \left[ LTRPN \right]_{tot}-\left[ LTRPNCa \right] \right)-k_{ltrpn}^{-}\left[ LTRPNCa \right]\#\left( \right) \end{aligned}$$

$$\begin{aligned} \frac{d\left[ HTRPNCa \right]}{dt}=k_{htrpn}^{+}\left[ Ca^{2+} \right]_{i}\left( \left[ HTRPN \right]_{tot}-\left[ HTRPNCa \right] \right)-k_{htrpn}^{-}\left[ HTRPNCa \right]\#\left( \right) \end{aligned}$$

$$\frac{dO_{1}}{dt}=-\left( k_{a}^{-}O_{1}+k_{b}^{+}\left[ Ca^{2+} \right]_{ss}^{m}O_{1}+k_{c}^{+}O_{1} \right)+$$

$$\begin{aligned} k_{a}^{+}\left[ Ca^{2+} \right]_{ss}^{n}P_{C1}+k_{b}^{-}O_{2}+k_{c}^{-}P_{C2}\#\left( \right) \end{aligned}$$

$$\begin{aligned} \frac{dO_{2}}{dt}=k_{b}^{+}\left[ Ca^{2+} \right]_{ss}^{m}O_{1}-k_{b}^{-}O_{2}\#\left( \right) \end{aligned}$$

$$\begin{aligned} \frac{dP_{C2}}{dt}=k_{c}^{+}O_{1}-k_{c}^{-}P_{C2}\#\left( \right) \end{aligned}$$

$$\begin{aligned} P_{C1}=1-\left( P_{C2}+O_{1}+O_{2} \right)\#\left( \right) \end{aligned}$$

$$\begin{aligned} \frac{dO}{dt}=-\left( 4\beta O+\gamma O+\alpha C_{4} \right)+K_{pcb}I_{1}+0.001\left( \alpha I_{2}-K_{pcf}O \right)\#\left( \right) \end{aligned}$$

$$\begin{aligned} \frac{dC_{2}}{dt}=-\left( \beta C_{2}+3\alpha C_{2} \right)+4\alpha C_{1}+2\beta C_{3}\#\left( \right) \end{aligned}$$

$$\begin{aligned} \frac{dC_{3}}{dt}=-\left( 2\beta C_{3}+2\alpha C_{3} \right)+3\alpha C_{2}+3\beta C_{4}\#\left( \right) \end{aligned}$$

$$\frac{dC_{4}}{dt}=-\left( 3\beta C_{4}+\alpha C_{4}+\gamma K_{pcf}C_{4} \right)+2\alpha C_{3}+4\beta O+$$

$$\begin{aligned} 0.01\left( 4K_{pcb}\beta I_{1}-\alpha\gamma C_{4} \right)+0.002\left( 4\beta I_{2}-K_{pcf}C_{4} \right)+4\beta K_{pcb}I_{3}\#\left( \right) \end{aligned}$$

$$\frac{dI_{1}}{dt}=-K_{pcb}I_{1}+\gamma O+0.001\left( \alpha I_{3}-K_{pcf}I_{1} \right)+$$

$$\begin{aligned} 0.01\left( \alpha\gamma C_{4}-4\beta K_{pcf}I_{1} \right)\#\left( \right) \end{aligned}$$

$$\frac{dI_{2}}{dt}=-\gamma I_{2}+0.001\left( K_{pcf}O-\alpha I_{2} \right)+K_{pcb}I_{3}+$$

$$\begin{aligned} 0.002\left( K_{pcf}C_{4}-4\beta I_{2} \right)\#\left( \right) \end{aligned}$$

$$\frac{dI_{3}}{dt}=-\left( 4\beta K_{pcb}I_{3}+K_{pcb}I_{3} \right)+0.001\left( K_{pcf}I_{1}-\alpha I_{3} \right)+$$

$$\begin{aligned} \gamma I_{2}+\gamma K_{pcf}C_{4}\#\left( \right) \end{aligned}$$

$$\begin{aligned} \alpha=\frac{0.4e^{0.1\left( V+12.0 \right)}\left[ 1.0-0.75 e^{-0.0025\left( V+20 \right)^{2}}+0.7 e^{-0.1\left( V+40 \right)^{2}} \right]}{1+0.12 e^{0.1\left( V+12 \right)}}\#\left( \right) \end{aligned}$$

$$\begin{aligned} \beta=0.05e^{-\left( V+12 \right)/13}\#\left( \right) \end{aligned}$$

$$\begin{aligned} \gamma=\frac{K_{pc,max}\left[ Ca^{2+} \right]_{ss}}{K_{pc,half}+\left[ Ca^{2+} \right]_{ss}}\#\left( \right) \end{aligned}$$

$$\begin{aligned} K_{pcf}=13\left[ 1-e^{-0.01\left( V+14.5 \right)^{2}} \right]\#\left( \right) \end{aligned}$$

$$\begin{aligned} I_{pCa}=I_{pCa}^{max}\frac{\left[ Ca^{2+} \right]_{i}^{2}}{K_{m,pCa}^{2}+\left[ Ca^{2+} \right]_{i}^{2}}\#\left( \right) \end{aligned}$$

$$\begin{aligned} I_{Cab}=G_{Cab}\left( V-E_{CaN} \right)\#\left( \right) \end{aligned}$$

$$I_{NaCa}=k_{NaCa}\left( \frac{1}{K_{m,Na}^{3}+\left[ Na^{+} \right]_{o}^{3}} \right)\left( \frac{1}{K_{m,Ca}+\left[ Ca^{2+} \right]_{o}} \right)$$

$$\begin{aligned} \cdot\left( \frac{1}{1+k_{sat}e^{\left( \eta-1 \right)VF/RT}} \right)\times\{e^{\eta VF/RT}\left[ Na^{+} \right]_{i}^{3}\left[ Ca^{2+} \right]_{o}-e^{\left( \eta-1 \right)VF/RT}\left[ Na^{+} \right]_{o}^{3}\left[ Ca^{2+} \right]_{i}\}\#\left( \right) \end{aligned}$$

$$\begin{aligned} I_{Cl,Ca}=G_{Cl,Ca}O_{Cl,Ca}\left( V-E_{Cl} \right)\frac{\left[ Ca^{2+} \right]_{i}}{\left[ Ca^{2+} \right]_{i}+K_{m,Cl}}\#\left( \right) \end{aligned}$$

$$\begin{aligned} O_{Cl,Ca}=\frac{0.2}{1+e^{-\left( V-46.7 \right)/7.8}}\#\left( \right) \end{aligned}$$

$$\begin{aligned} g_{KATP}=i_{KATP_{on}}\frac{0.000193}{N_{area}}\#\left( \right) \end{aligned}$$

$$\begin{aligned} p_{ATP}=\frac{1}{1+\left( \frac{\left[ ATP \right]_{i}}{k_{ATP}} \right)^{h_{ATP}}}\#\left( \right) \end{aligned}$$

$$\begin{aligned} G\bar{K}T=g_{KATP}p_{ATP}\left( \frac{Ko}{4} \right)^{n_{ATP}}\#\left( \right) \end{aligned}$$

$$\begin{aligned} I_{KATP}=G\bar{K}T\left( V-E_{K} \right)\#\left( \right) \end{aligned}$$

$$\begin{aligned} t_{s}=t-t_{start}\#\left( \right) \end{aligned}$$

$$\begin{aligned} r_{star}=r_{0}+pulse\left[ heav\left( mod\left( t_{s},period \right)-t_{f} \right)-heav\left( mod\left( t_{s},period \right)-\left( t_{f}+t_{p} \right) \right) \right]\#\left( \right) \end{aligned}$$

Appendix A

**Table A1.** List of parameters in the model

| **Parameter** | **Description** | **Value** |
| --- | --- | --- |
| $A_{cap}$ | Capacitive membrane area | $1.534\times{10}^{-4}cm^{2}$ |
| $V_{myo}$ | Myoplasmic volume | $2.584\times{10}^{-5}\mu l$ |
| $V_{JSR}$ | Junctional SR volume | $1.2\times{10}^{-7}\mu l$ |
| $V_{NSR}$ | Network SR volume | $2.098\times{10}^{-6}\mu l$ |
| $V_{ss}$ | Subspace volume | $1.485\times{10}^{-9}\mu l$ |
| $\left[ Na^{+} \right]_{i}$ | Myoplasmic $Na^{+}$ concentration | $14237.1 \mu M$ |
| $\left[ K^{+} \right]_{i}$ | Myoplasmic $K^{+}$concentration | $143720.0 \mu M$ |
| $\left[ Ca^{2+} \right]_{i}$ | Myoplasmic $Ca^{2+}$ concentration | $0.115001 \mu M$ |
| $\left[ K^{+} \right]_{o}$ | Extracellular $K^{+}$ concentration | $5400.0 \mu M$ |
| $\left[ Na^{+} \right]_{o}$ | Extracellular $Na^{+}$ concentration | $140000.0 \mu M$ |
| $\left[ Ca^{2+} \right]_{o}$ | Myoplasmic $Ca^{2+}$ concentration | $1800.0 \mu M$ |
| $\left[ Ca^{2+} \right]_{ss}$ | Subspace SR $Ca^{2+}$ concentration | $0.115001 \mu M$ |
| $\left[ Ca^{2+} \right]_{JSR}$ | JSR $Ca^{2+}$ concentration | $1299.50 \mu M$ |
| $\left[ Ca^{2+} \right]_{NSR}$ | NSR $Ca^{2+}$ concentration | $1299.50 \mu M$ |
| $\left[ LTRPNCa \right]$ | Concentration $Ca^{2+}$ bound low-affinity troponin-binding sites | $11.2684 \mu M$ |
| $\left[ HTRPNCa \right]$ | Concentration $Ca^{2+}$ bound high-affinity troponin-binding sites | $125.290 \mu M$ |
| $\left[ LTRPN \right]_{tot}$ | Total myoplasmic troponin low-affinity site concentration | $70.0 \mu M$ |
| $\left[ HTRPN \right]_{tot}$ | Total myoplasmic troponin high-affinity site concentration | $140.0 \mu M$ |
| $C_{m}$ | Specific membrane capacitance | $1.0 \mu F/cm^{2}$ |
| $F$ | Faraday constant | $96.5 C/mmol$ |
| $T$ | Absolute temperature | $298 K$ |
| $R$ | Ideal gas constant | $8.314 J\cdot mol^{-1}\cdot K^{-1}$ |
| $E_{Ca,L}$ | Reversal potential for L-type $Ca^{2+}$ channel | $63.0 mV$ |
| $E_{Cl}$ | Reversal potential for $Ca^{2+}$-activated $Cl^{-}$ current | $-40.0 mV$ |
| $k_{NaCa}$ | Scaling factor of $Na^{+}/Ca^{2+}$ exchage | $992.8 pA/pF$ |
| $K_{m,Na}$ | $Na^{+}$ half-saturation constant for $Na^{+}/Ca^{2+}$ exchange | $87500.0 \mu M$ |
| $K_{m,Ca}$ | $Ca^{2+}$ half-saturation constant for $Na^{+}/Ca^{2+}$ exchange | $1380 \mu M$ |
| $k_{sat}$ | $Na^{+}/Ca^{2+}$exchange saturation factor at very negative potentials | 0.1 |
| $\eta$ | Controls voltage dependence of $Na^{+}/Ca^{2+}$ exchange | 0.35 |
| $I_{NaK}^{max}$ | Maximum $Na^{+}/K^{+}$ exchange current | $0.88 pA/pF$ |
| $K_{m,Nai}$ | $Na^{+}$ half-saturation constant for $Na^{+}/K^{+}$exchange current | $21.0 mM$ |
| $K_{m,Ko}$ | $K^{+}$ half-saturation constant for $Na^{+}/K^{+}$ exchange current | $1.5 mM$ |
| $I_{pCa}^{max}$ | Maximum $Ca^{2+}$ pump current | $1.0 pA/pF$ |
| $K_{m,pCa}$ | $Ca^{2+}$ half-saturation constant for $Ca^{2+}$ pump current | $0.5 \mu M$ |
| $G_{Na}$ | Maximum fast $Na^{+}$ current conductance | $13 mS/\mu F$ |
| $G_{CaL}$ | Specific maximum conductivity for L-type $Ca^{2+}$ channel | $1.51729 mS/\mu F$ |
| $G_{Ks}$ | Maximum slow delayed-rectifier $K^{+}$current conductance | $0.000575 mS/\mu F$ |
| $G_{Kr}$ | Maximum rapid delayed-rectifier $K^{+}$ current conductance | $0.0078 mS/\mu F$ |
| $G_{Kur}$ | Maximum ultrarapidly delayed-rectifier $K^{+}$ current conductance(apex) | $0.0016 mS/\mu F$ |
| $G_{Cl,Ca}$ | Maximum $Ca^{2+}$-activated $Cl^{-}$ current conductance | $10.0 mS/\mu F$ |
| $G_{Kss}$ | Maximum noninactivating steady-state $K^{+}$ current conductance (apex) | $0.05 mS/\mu F$ |
| $G_{Cab}$ | Maximum background $Ca^{2+}$ current conductance | $3.67\times{10}^{-4} mS/\mu F$ |
| $G_{Nab}$ | Maximum background $Na^{+}$ current conductance | $0.0026 mS/\mu F$ |
| $G_{Kto,f}$ | Maximum transient outward $K^{+}$ current conductance (apex) | $0.4067 mS/\mu F$ |
| $G_{Kto,s}$ | Maximum transient outward $K^{+}$ current conductance (apex) | $0.01 mS/\mu F$ |
| $K_{pc,max}$ | Maximum time constant for $Ca^{2+}$-induced inactivation | $0.23324 ms^{-1}$ |
| $K_{pc,half}$ | Half-saturation constant for $Ca^{2+}$-induced inactivation | $20.0 \mu M$ |
| $K_{pcb}$ | Voltage-insensitive rate constant for inactivation | $5.0\times{10}^{-4} ms^{-1}$ |
| $I_{CaL,max}$ | Normalization constant for L-type $Ca^{2+}$ current | $7.0 pA/pF$ |
| $\nu_{1}$ | Maximum RyR channel $Ca^{2+}$ permeability | $4.5 ms^{-1}$ |
| $\nu_{2}$ | $Ca^{2+}$ leak rate constant from the NSR | $1.7\times{10}^{-5} ms^{-1}$ |
| $\nu_{3}$ | SR $Ca^{2+}$-ATPase maximum pump rate | $0.45 \mu M/ms$ |
| $K_{m,up}$ | Half-saturation constant for SR $Ca^{2+}$-ATPase pump | $0.5 \mu M$ |
| $\tau_{xfer}$ | Time constant for transfer from subspace to myoplasm | $8.0 ms$ |
| $\tau_{tr}$ | Time constant for transfer from NSR to JSR | $20.0 ms$ |
| $k_{a}^{+}$ | RyR $P_{C1}$ - $P_{O1}$ rate constant | $0.006075 \mu M^{-4}/ms$ |
| $k_{b}^{+}$ | RyR $P_{O1}$ - $P_{O2}$ rate constant | $0.00405 \mu M^{-3}/ms$ |
| $k_{a}^{-}$ | RyR $P_{O1}$ - $P_{C1}$ rate constant | $0.07125 ms^{-1}$ |
| $k_{b}^{-}$ | RyR $P_{O2}$ - $P_{O1}$ rate constant | $0.965 ms^{-1}$ |
| $k_{c}^{-}$ | RyR $P_{C2}$ - $P_{O1}$ rate constant | $8.0\times{10}^{-4} ms^{-1}$ |
| $k_{c}^{+}$ | RyR $P_{O1}$ - $P_{C2}$ rate constant | $0.0090 ms^{-1}$ |
| $n$ | RyR $Ca^{2+}$ cooperativity parameter $P_{C1}$ - $P_{O1}$ | 4.0 |
| $m$ | RyR $Ca^{2+}$ cooperativity parameter $P_{O1}$ - $P_{O2}$ | 3.0 |
| $\left[ LTRPN \right]_{tot}$ | Total myoplasmic troponin low-affinity site concentration | $70.0 \mu M$ |
| $\left[ HTRPN \right]_{tot}$ | Total myoplasmic troponin high-affinity site concentration | $140.0 \mu M$ |
| $\left[ CSQN \right]_{tot}$ | Total junctional SR calsequestrin concentration | $15000.0 \mu M$ |
| $\left[ CMDN \right]_{tot}$ | Total myoplasmic calmodulin concentration | $50.0 \mu M$ |
| $\left[ LTRPNCa \right]_{tot}$ | Concentration $Ca^{2+}$ bound low-affinity troponin-binding sites | $11.2684 \mu M$ |
| $\left[ HTRPNCa \right]_{tot}$ | Concentration $Ca^{2+}$ bound high-affinity troponin-binding sites | $125.29 \mu M$ |
| $k_{ltrpn}^{+}$ | $Ca^{2+}$ on rate constant for troponin low affinity sites | $0.0327 \mu M^{-1}/ms$ |
| $k_{htrpn}^{+}$ | $Ca^{2+}$ on rate constant for troponin high affinity sites | $0.00237 \mu M^{-1}/ms$ |
| $k_{ltrpn}^{-}$ | $Ca^{2+}$ off rate constant for troponin low affinity sites | $0.196 ms^{-1}$ |
| $k_{htrpn}^{-}$ | $Ca^{2+}$ off rate constant for troponin high affinity sites | $3.2\times{10}^{-5} ms^{-1}$ |
| $K_{m}^{CMDN}$ | $Ca^{2+}$ half-saturation constant for calmodulin | $0.238 \mu M$ |
| $K_{m}^{CSQN}$ | $Ca^{2+}$ half-saturation constant for calsequestrin | $800.0 \mu M$ |
| $k_{f}$ | Rate constant for rapid delayed-rectifier $K^{+}$ current | $0.023761 ms^{-1}$ |
| $k_{b}$ | Rate constant for rapid delayed-rectifier $K^{+}$ current | $0.036778 ms^{-1}$ |
| $K_{m,Cl}$ | Half-saturation constant for $Ca^{2+}$-activated $Cl^{-}$ current | $10.0 \mu M$ |
| $O$ | L-type $Ca^{2+}$ channel conducting state | $9.30308\times{10}^{-19}$ |
| $O_{1}$ | Fraction of RyR channels in state $O_{1}$ | $1.49102\times{10}^{-5}$ |
| $O_{2}$ | Fraction of RyR channels in state $O_{2}$ | $9.51726\times{10}^{-11}$ |
| $C_{1}$ | L-type $Ca^{2+}$ channel closed state | 0.999876 |
| $C_{2}$ | L-type $Ca^{2+}$ channel closed state | $1.24216\times{10}^{-4}$ |
| $C_{3}$ | L-type $Ca^{2+}$ channel closed state | $5.78679\times{10}^{-9}$ |
| $C_{4}$ | L-type $Ca^{2+}$ channel closed state | $1.19816\times{10}^{-13}$ |
| $I_{1}$ | L-type $Ca^{2+}$ channel inactivated state | $4.97023\times{10}^{-19}$ |
| $I_{2}$ | L-type $Ca^{2+}$ channel inactivated state | $3.45847\times{10}^{-14}$ |
| $I_{3}$ | L-type $Ca^{2+}$ channel inactivated state | $1.85106\times{10}^{-14}$ |
| $P_{C1}$ | Fraction of RyR channels in state $P_{C1}$ | 0.999817 |
| $P_{C2}$ | Fraction of RyR channels in state $P_{C2}$ | $1.6774\times{10}^{-4}$ |
| $P_{RyR}$ | RyR modulation factor | 0.0 |
| $C_{Na1}$ | Closed state of fast $Na^{+}$ channel | $2.79132\times{10}^{-4}$ |
| $C_{Na2}$ | Closed state of fast $Na^{+}$ channel | 0.020752 |
| $C_{Na3}$ | Closed state of fast $Na^{+}$ channel | 0.624646 |
| $O_{Na}$ | Closed state of fast $Na^{+}$ channel | $7.13483\times{10}^{-7}$ |
| $IF_{Na}$ | Fast inactivated state of fast $Na^{+}$ channel | $1.53176\times{10}^{-4}$ |
| $I1_{Na}$ | Slow inactivated state 1 of fast $Na^{+}$ channel | $6.73345\times{10}^{-7}$ |
| $I2_{Na}$ | Slow inactivated state 2 of fast $Na^{+}$ channel | $1.55787\times{10}^{-9}$ |
| $IC_{Na2}$ | Closed-inactivated state of fast $Na^{+}$ channel | 0.0113879 |
| $IC_{Na3}$ | Closed-inactivated state of fast $Na^{+}$ channel | 0.34278 |
| $a_{to,f}$ | Gating variable for transient outward $K^{+}$ current | 0.00265563 |
| $i_{to,f}$ | Gating variable for transient outward $K^{+}$ current | 0.999977 |
| $a_{to,s}$ | Gating variable for transient outward $K^{+}$ current | $4.17069\times{10}^{-4}$ |
| $i_{to,s}$ | Gating variable for transient outward $K^{+}$ current | 0.998543 |
| $n_{Ks}$ | Gating variable for slow delayed-rectifier $K^{+}$ current | $2.62753\times{10}^{-4}$ |
| $a_{ur}$ | Gating variable for ultra rapidly activating delayed-rectifier $K^{+}$ current | $4.17069\times{10}^{-4}$ |
| $i_{ur}$ | Gating variable for ultra rapidly activating delayed-rectifier $K^{+}$ current | 0.998543 |
| $a_{Kss}$ | Gating variable for noninactivating steady-state $K^{+}$ current | $4.17069\times{10}^{-4}$ |
| $i_{Kss}$ | Gating variable for noninactivating steady-state $K^{+}$ current | 1 |
| $C_{K0}$ | mERG channel closed state | 0.998159 |
| $C_{K1}$ | mERG channel closed state | $9.92513\times{10}^{-4}$ |
| $C_{K2}$ | mERG channel closed state | $6.41229\times{10}^{-4}$ |
| $O_{K}$ | mERG channel open state | $1.75298\times{10}^{-4}$ |
| $I_{K}$ | mERG channel inactivated state | $3.19129\times{10}^{-5}$ |
| $i_{K_{ATP_{on}}}$ | current passing through the open KATP channels | 1 |
| $n_{\text{ATP}}$ | activation threshold of ATP molecules on the KATP channel | 0.24 |
| $h_{ATP}$ | enhanced effect of ATP on KATP channel opening | 2 |
| $k_{ATP}$ | ATP sensitivity constant of the KATP channel | 0.00025 |
| $N_{area}$ | number of ion channels per unit area | $5\times{10}^{-5}$ |
| $\left[ ATP \right]_{i}$ | intracellular concentration of ATP | $3 mM$ |
| $V$ | Membrane potential | $-82.4202 mV$ |
| $t$ | Time | $0.0 ms$ |
| $r_{0}$ | Initial amplitude | $1 ms$ |
| $period$ | Stimulation cycle | $300 ms$ |
| $pulse$ | Stimulation amplitude | $10 pA/pF$ |
| $t_{start}$ | Initial time lapse | $10 ms$ |

Appendix B

The following script was used for STFT of calcium signals or intensity signals from a kymograph.

This script reads an Excel file, extracts the data column, and then calculates and plots a spectrogram using the Short-Time Fourier Transform (STFT).

import pandas as pd

import numpy as np

import matplotlib.pyplot as plt

from scipy import signal

*# Read Excel file*

df = pd.read_excel('your file')

*# Get the column data*

data = df.iloc[:, 2].values

fs = 1 / 0.1312 *# Sampling frequency*

nperseg = 300 *# Each window length is 256 points*

noverlap = 295 *# Windows overlap by half*

nfft = 300 *# Use 1024-point Fast Fourier Transform*

*# Calculate spectrogram*

freqs, times, spectrogram = signal.spectrogram(data, fs=fs, nperseg=nperseg, noverlap=noverlap, nfft=nfft)

plt.rcParams['figure.dpi'] = 300 *# Resolution*

plt.rcParams['font.family'] = 'Arial' *# Font*

plt.rcParams['font.size'] = 10 *# Font size*

plt.rcParams['axes.spines.top'] = False *# Do not plot top border*

plt.rcParams['axes.spines.right'] = False *# Do not plot right border*

spectrogram[freqs > 1.5] = 0

*# Plot spectrogram*

plt.pcolormesh(times, freqs, spectrogram, cmap='coolwarm')

plt.xlabel('Time [sec]')

plt.ylabel('Frequency [Hz]')

plt.ylim([0, 4]) *# Set y-axis range*

plt.colorbar()

plt.axis('off')

plt.show()
